# Splicing inhibition enhances the antitumor immune response through increased tumor antigen presentation and altered MHC-I immunopeptidome

**DOI:** 10.1101/512681

**Authors:** Alison Pierson, Romain Darrigrand, Marine Rouillon, Mathilde Boulpicante, Zafiarisoa Dolor Renko, Camille Garcia, Michael Ghosh, Marie-Charlotte Laiguillon, Camille Lobry, Mouad Alami, Sébastien Apcher

**Author notes:** These authors contributed equally to this work (Alison Pierson, Romain Darrigrand). Corresponding Author: Sébastien Apcher, Gustave Roussy, University Paris Sud, University Paris Saclay, INSERM U1015, 114 rue Edouard Vaillant,94805 Villejuif, France.

## Abstract

The success of cancer immunotherapy relies on the induction of an immunoprotective response targeting tumor antigens (TAs) presented by tumor cells on MHC class I molecules. Alternative translation events emerged as a rich source of TAs and generate the so-called Pioneer Translation Products (PTPs), which are peptides generated from unspliced mRNA. We demonstrated in vitro and in vivo that the splicing inhibitor isoginkgetin and a derived water-soluble and less toxic molecule, IP2, act at the production stage of the PTPs. We showed that IP2 increases PTP-derived antigen presentation in cancer cells *in vitro* and decreases tumor growth *in vivo* in an immune-dependent manner. Furthermore, IP2 treatment induces a long-lasting antitumor response. Finally, we observed that the epitope repertoire displayed on MHC-I molecules is altered upon treatment with IP2 with the modulation of pre-existing peptides and the emergence of novel antigens derived from both coding and allegedly non-coding sequences.

**Significance:** IP2 is a new efficient “first in class” immunomodulator of the MHC I presentation pathway. IP2 reduces the growth of sarcoma MCA205 and melanoma B16F10 tumors bearing the PTP-derived SL8 epitope and significantly extends mice survival. IP2 treatment reshape the cancer cell MHC-I immunopeptidome. These findings add to the understanding of the role of the splicing machinery in antigen production and presentation and identify the spliceosome as a druggable target to enhance cancer immunosurveillance.

---

All nucleated cells of jawed vertebrates present antigenic peptides (APs) at their surface through the major histocompatibility complex class I (MHC-I) presentation pathway (1). APs are 8- to 12- amino acid long and reflect the inherent metabolic cellular activity. Initial clinical trials using vaccines targeting tumor antigens (TA) have not met their expectations. The main failures have been associated with immunosuppressive mechanisms and with a suboptimal choice of antigens (2,3). One of the important events that drives tumor immunoselection and that is correlated with a poor prognosis is the loss or the downregulation of MHC-I antigen presentation by tumor cells (4,5). The latter can escape cytotoxic T lymphocyte (CTL) and natural killer cell recognition thanks to defects in components of the MHC-I pathway (6). Moreover, along with the overall decrease in MHC-I antigen presentation, the very nature and the amount of antigens presented at the cell surface, namely the MHC-I immunopeptidome, is of critical importance for immune recognition. For example, the loss of expression at the tumor cell surface of a specific TA identified and targeted with immunotherapy, such as Her/neu in breast cancer or CEA in colon cancer, leads to immune evasion (7,8). To counteract this phenomenon, current strategies aim at enlarging the range of targeted cancer peptides and restoring MHC-I antigen presentation (9–12).

In order to understand the dynamic of the MHC-I immunopeptidome, we focused on the source of APs for the MHC-I presentation pathway. Exploring the concept of Defective Ribosomal Products (DRIPs) (13–16), we showed that one of the most important source of APs is a pioneer translation event that occurs on pre-mRNAs, before introns are spliced out, and independently of the translation of the corresponding full length proteins. Produced non-canonical peptides can therefore be derived from intronic sequences, 3’ or 5’ UTR regions as well as alternative reading frames. These polypeptides are described as Pioneer Translation Products (PTPs) (17,18). The discovery of PTPs emphasizes the existence of a nuclear translational mechanism of precursors-mRNAs that participates to the production of MHC-I peptides (MIPs) by generating suitable polypeptides for the MHC-I pathway. Moreover, PTPs play a role in the dynamic of cancer development. When inoculated into mice, it has been shown that cancer cells presenting PTP-derived antigens at their surface can be recognized by specific T-cells leading to tumor growth reduction. Besides, purified PTPs containing a model epitope efficiently promote anti-cancer immune response when injected as a peptide vaccine into mice (19).

Precursor-mRNA (pre-mRNA) splicing is catalyzed in the nucleus by the spliceosome, a conserved and dynamic multi-RNA/protein complex composed of five small nuclear RNAs (snRNAs) in interaction with over 180 proteins (20). A growing number of studies report that the deregulation of the spliceosome complex entails aberrant splicing patterns in many cancers contributing to abnormal tumor cell proliferation and progression (21–24). Furthermore, cancer cells have different intracellular mechanisms to shape the pool of peptides presented on MHC-I molecules at their surface, leading to the reduction of their immunogenicity and allowing them to escape from T-cell recognition (6,25). In a recent study, we observed that splicing inhibition positively modulate the surface presentation of a PTP-derived model antigen in healthy cells treating with isoginkgetin (18). The latter molecule has been reported to inhibit the spliceosome during the early stages of its assembly (26). From these observations, the link between splicing inhibition and antigen presentation in cancer cells became of particular interest for us and splicing inhibitors appeared as one potential strategy to render tumor cells more visible to CTL. Since 2011, recurrent spliceosome mutations have been reported in several cancers. Among strategies developed to render tumor cells more visible to CTL, a novel class of anticancer molecules has been of particular interest for us: the splicing inhibitors (21). Several natural products and their synthetic analogues have been reported to inhibit the spliceosome, including pladienolides B and D, spliceostatin A, FR901464, E7107, isoginkgetin and madrasin. Beyond their previously reported cytotoxic activities (27,28), we aimed at testing their effect on PTP production and antigen presentation.

Here we show that the biflavonoid isoginkgetin and its water-soluble derivative IP2 we generated enhance the presentation of PTP-derived antigens in human and mouse cancer cells *in-vitro*. In addition, IP2 induces a long-lasting anti-cancer immune response *in vivo*. Finally, IP2 was shown to re shape the MHC-I immunopeptidome, leading to a new peptide representation at the cell surface. Our findings suggest a new mechanism of action of those splicing inhibitors that modulate the presentation of MIPs at the cancer cell surface by enhancing the presentation of PTP-derived antigens and potentiate the antitumor immune response.

## Results

### Splicing inhibition increases the presentation of conventional and non-conventional antigens in cancer cells

To improve antigenicity and immune recognition of cancer cells, we determined whether isoginkgetin was able to modulate positively the expression and the presentation of tumor associated PTP-derived antigens at the surface of cancer cells. For that purpose, the human melanoma cell line A375, the human lung cancer cell line A549 and the normal human fibroblast lung cell line MRC5 transiently expressing the mouse MHC-I H-2K^b^ molecule and the intron-derived SL8 epitope within the β-Globin gene construct (Globin-SL8-intron) were treated at different doses of isoginkgetin for 18h. The overall expression of the MHC-I H-2K^b^ molecules at the cell surface upon treatment was assessed by B3Z activation. Treatment with isoginkgetin increases intron-derived SL8-antigen presentation in the three cell types, in a dose dependent manner (Fig. 1A). In parallel, the same experiment was performed on the mouse sarcoma MCA205 and mouse melanoma B16F10 cell lines that were transiently expressing the Globin-SL8-intron construct. In accordance with the previous results, isoginkgetin elicited an increase in the intron-derived SL8 antigen presentation, in a dose dependent manner (Fig. 1B). To investigate further the impact of isoginkgetin on PTP presentation, MCA205 and B16F10 cell lines transiently expressing the exon-derived SL8 epitope within the β-Globin gene construct (Globin-SL8-exon) or the splicing independent Ova-derived SL8 were treated with increasing doses of the compound. We observed that isoginkgetin increases splicing dependent but not splicing independent SL8 presentation in a dose dependent manner (Figs. 1C and 1D). Those results show that isoginkgetin-induced PTP-presentation is mediated through altered splicing events. This suggests an action of isoginkgetin during the production stage of PTPs and not downstream in the MHC-I antigen presentation pathway.

**Figure 1:**
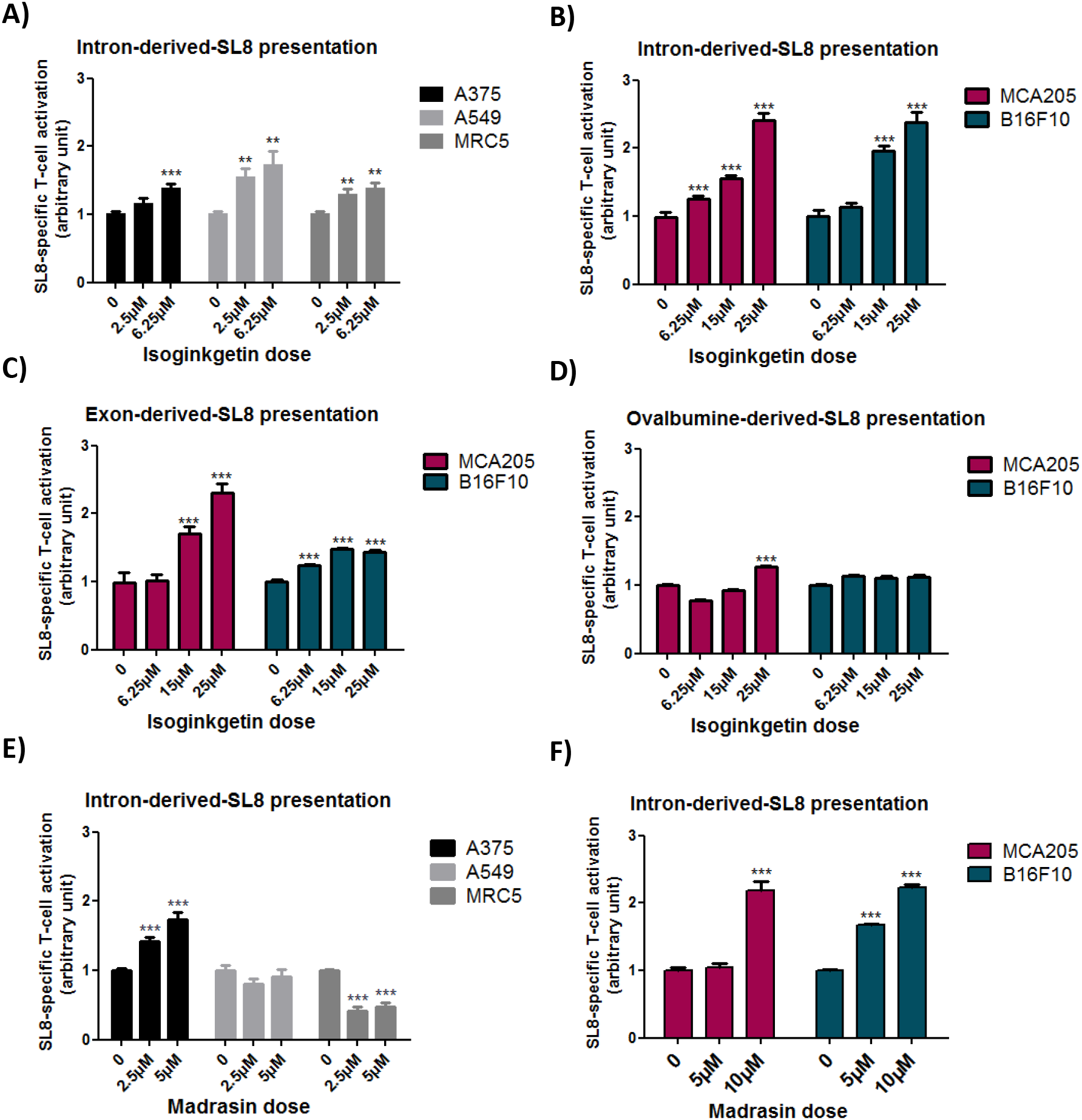
Splicing inhibition increases exon- and intron-derived antigen presentation in cancer cells. B3Z SL8-specific T-cell activation after co-culture with **(A)** human melanoma A375, human lung carcinoma A549 or normal human lung fibroblast MRC5 cell lines, all transiently expressing the mouse H2-K^b^ molecules and the intron-derived SL8 epitope and treated with 2.5μM or 6.25 μM of isoginkgetin for 18 hours or **(B)** mouse sarcoma MCA205 and mouse melanoma B16F10 cell lines both transiently expressing the intron-derived SL8 epitope and treated with 6.25μM, 15μM or 25μM of isoginkgetin for 18 hours. B3Z SL8-specific T-cell activation after co-culture with mouse MCA205 or mouse B16F10 cell lines both transiently expressing **(C)** the exon-derived SL8 epitope or **(D)** the Ova cDNA construct and treated with 6.25μM, 15μM or 25μM of isoginkgetin for 18 hours. B3Z SL8-specific T-cell activation after co-culture with **(E)** human melanoma A375, human lung carcinoma A549 or normal human lung fibroblast MRC5 cell lines, all transiently expressing the mouse H2-K^b^ molecules and the intron-derived SL8 epitope and treated with 2.5μM or 5μM of madrasin for 18 hours or **(F)** mouse sarcoma MCA205 and mouse melanoma B16F10 cell lines both transiently expressing the intron-derived SL8 epitope and treated with 5μM or 10 μM of madrasin for 18 hours. Free soluble SL8 peptide was added to ensure that T-cell activation assays were carried out in non-saturated conditions. Unspecific B3Z T-cell activation was taking into account in the results. Each graph is one representative of at least four independent experiments. Data are given as mean ± SEM. *P<0.05, **P<0.01, ***P<0.001 (unpaired student t test).

Moreover, we assessed the ability of a second splicing inhibitor, the madrasin, to increase PTP-derived antigen presentation in cancer cells. Interestingly, as isoginkgetin did, madrasin increased intron-derived SL8 presentation in the MCA205 and B16F10 mouse cell lines and in the A375 human melanoma cell line (Figs. 1E and 1F). However madrasin was shown to be more toxic than isoginkgetin at the effective doses (Supplementary Fig. S1). In contrast, when treated with the chemotherapeutics gemcitabine or cyclophosphamide, the same cell lines that express the SL8 epitope from an intron did not show any significant modulation of the antigen expression (Supplementary Fig. S2), apart from MCA205 and A549 cells treated with gemcitabine. Taken together, those results suggest a specificity of action of the splicing inhibitor family on antigen presentation which is not shared with molecules with other activities.

Finally, we observed that the expression of the MHC-I H-2K^b^ molecules at the cell surface varies upon treatment depending on the cell type (Supplementary Fig. S3) but is not correlated to the effect of the drugs on SL8 antigen presentation.

Overall, these results show that the natural product isoginkgetin acts as a enhancer of the PTP-derived antigen presentation in cancer cells independently of the epitope setting, i.e. in exonic or in intronic sequences, and independently of the cell type. Moreover, the effects of both isoginkgetin and madrasin treatments shed further light on the importance of the splicing event for the production and the presentation of MHC-I antigens in cancer cells. Finally, they support the idea that pre-mRNAs are a source for antigen presentation when the spliced machinery is impaired.

### Isoginkgetin treatment slows down *in vivo* the growth of intron-derived SL8-expressing tumors

Antigen abundance at the cell surface has been demonstrated to be a key parameter in determining the magnitude of the CD8+ T cell response and hence in defining immunodominance (29). The SL8 peptide is highly immunogenic *in vivo*. Looking at SL8-specific T-cell activation *in vitro*, we observed an increase in the abundance of the SL8 expression at the cancer cell surface after splicing inhibition. In order to test this hypothesis *in vivo*, we first looked at the impact of isoginkgetin treatment on the growth of tumors expressing the intron-derived SL8 peptide. For that purpose, MCA205 sarcoma cells and B16F10 melanoma cells stably expressing the Globin-SL8-intron construct were inoculated subcutaneously into mice. At days 5, 10 and 15 post tumor inoculation, the mice were injected intraperitoneally with isoginkgetin and the tumor growth was monitored (Fig. 2A). In mice bearing MCA205 globin-SL8-intron tumors, we observed a significant reduction of tumor size, over 60% at day 27 after challenge with 12 and 18mg/kg of isoginkgetin (Fig. 2B and Supplementary Fig. S4A, left panels). The impact of isoginkgetin treatment on B16F10 globin-SL8-intron tumor growth is lower with over 30% of tumor reduction at day 18 after challenge with 12 and 18mg/kg of isoginkgetin (Fig. 2B and Supplementary Fig. S4A, right panels). To assess the link between SL8 overexpression and tumor growth reduction *in vivo* after isoginkgetin treatment, we performed the same experiment in mice inoculated with either MCA205 or B16F10 wild type (WT) cells. No significant reduction of the growth of MCA205 WT (Fig. 2C and Supplementary Fig. S4B, left panels) and B16F10 WT (Fig. 2C and Supplementary Fig. S4B, right panels) was observed after treatment with 12 and 18mg/kg of isoginkgetin. Taken together these results show that isoginkgetin induced PTP presentation and tumor growth reduction is mediated through altered splicing events.

**Figure 2:**
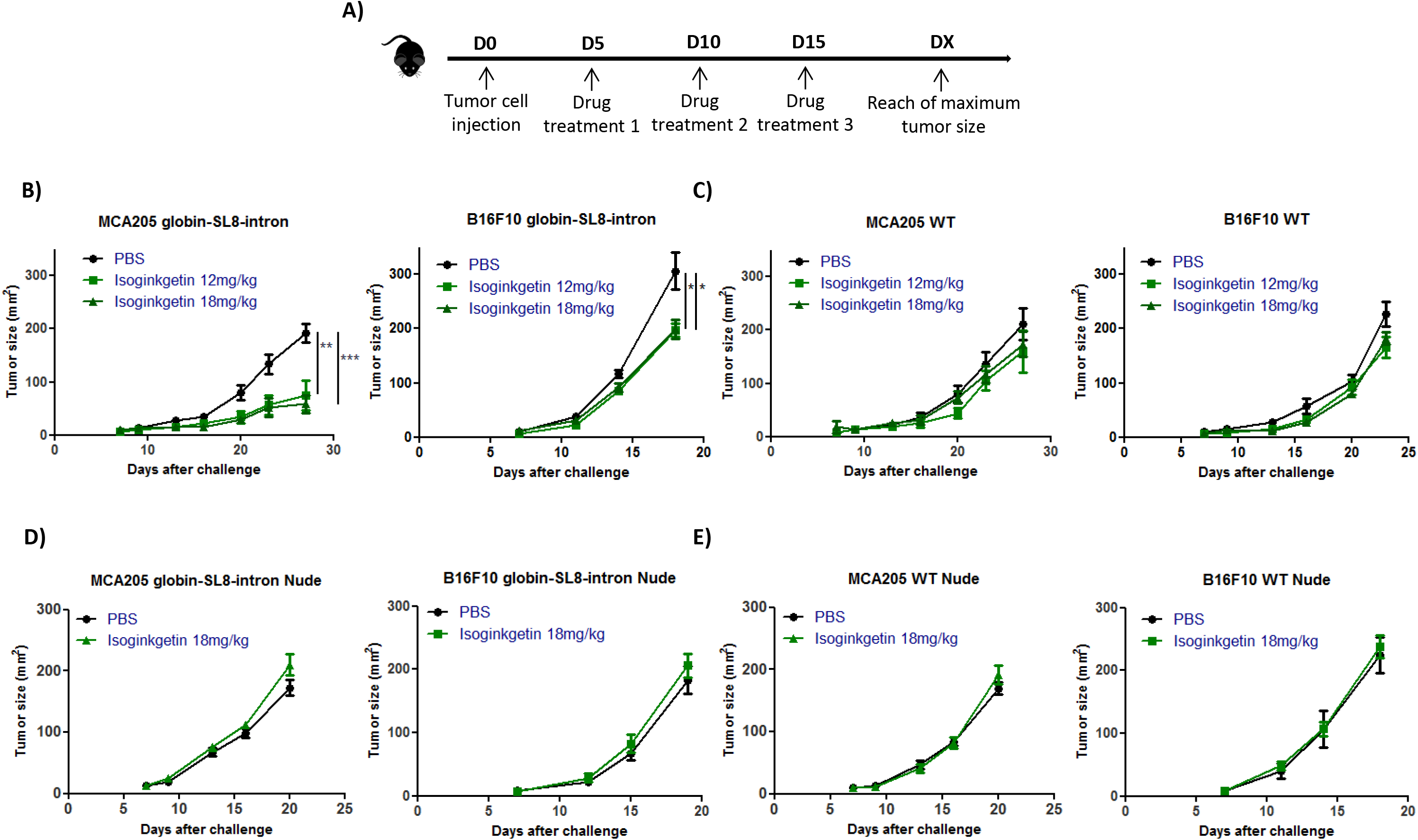
The splicing inhibitor isoginkgetin reduces the growth of tumor bearing intron-derived SL8 in an immune-dependent manner. **(A)** Experimental settings. Growth of **(B)** mouse sarcoma MCA205 (left panel) and mouse melanoma B16F10 (right panel) both stably expressing the globin-SL8-intron construct or **(C)** MCA205 Wild Type (WT) (left panel) and B16F10 WT cells (right panel) subcutaneously inoculated into the right flank of immunocompetent C57BL/6 mice thereafter treated intraperitoneally with 12mg/kg or 18mg/kg of isoginkgetin at day 5, 10 and 15 post tumor inoculation. Growth of **(D)** mouse sarcoma MCA205 (left panel) and mouse melanoma B16F10 (right panel) both stably expressing the globin-SL8-intron construct or **(E)** MCA205 Wild Type (WT) (left panel) and B16F10 WT cells (right panel) subcutaneously inoculated into the right flank of immunodeficient Nu/Nu mice thereafter treated intraperitoneally with 18mg/kg of isoginkgetin at day 5, 10 and 15 post tumor inoculation. Tumor size was assessed every 3 to 4 days until it reached the established ethical endpoints. Each line represents the tumor size in area (mm^2^) of at least 6 mice in each group. Data are given as mean ± SEM. *p<0.05, **p<0.01 (ANOVA with Tukey’s multiple comparison test comparing all groups).

We then assessed the requirement of the immune system for isoginkgetin to reduce tumor growth. Immunodeficient Nu/Nu nude mice were inoculated subcutaneously with either MCA205 or B16F10 cells stably expressing the Globin-SL8-intron construct or WT cells, and thereafter treated under the same conditions as previously described (Fig. 2A). No effect of isoginkgetin treatment was observed on the growth of each of the four tumor types (Figs. 2D-E and Supplementary Figs. S4C-D).

Overall, these results show that tumor growth slow down upon isoginkgetin treatment requires the presence of an active immune response *in vivo*, and suggest that the increase in the expression of an immunodominant epitope drives the antitumor immune response.

### The water-soluble IP2 and M2P2 compounds derived from the natural isoginkgetin product inhibit the splicing with reduced cytotoxicity

With the aim of transferring the isoginkgetin compound to the clinic, two of its intrinsic properties were limiting: its hydrophobicity which reduces its bioavailability and its cytotoxicity on both normal and tumor cells. Therefore, derivatives of the natural isoginkgetin product were synthesized and tested for their ability to inhibit the splicing machinery, to increase PTP-derived antigens *in vitro* as well as to reduce tumor growth *in vivo*.

From the commercial isoginkgetin (Fig. 3A), extracted from leaves of the maidenhair tree, *Ginko biloba*, the derivatives IP2 (Fig. 3B) and M2P2 (Fig. 3C) were synthesized. The synthesis route of each compound is provided in Figure S5. Briefly, the synthesis of IP2 (referred as compound 2 in the schematic) was accomplished by the phosphorylation of isoginkgetin employing *in situ* formation of diethylchlorophosphite to provide compound 1. Further cleavage of the ethyl ester protective groups with iodotrimethylsilane afforded the phosphoric acid intermediate, which was immediately treated with sodium hydroxide to complete a practical route to the disodium phosphate prodrug. For the synthesis of the M2P2 molecule, the remaining two phenol groups of compound 1 were alkylated using methyl iodide to furnish compound 3. Treatment of the latter under similar conditions to prepare compound 2 from compound 1 gave the disodium phosphate prodrug 4 or M2P2, whereas its reaction under basic conditions provided compound 5.

**Figure 3:**
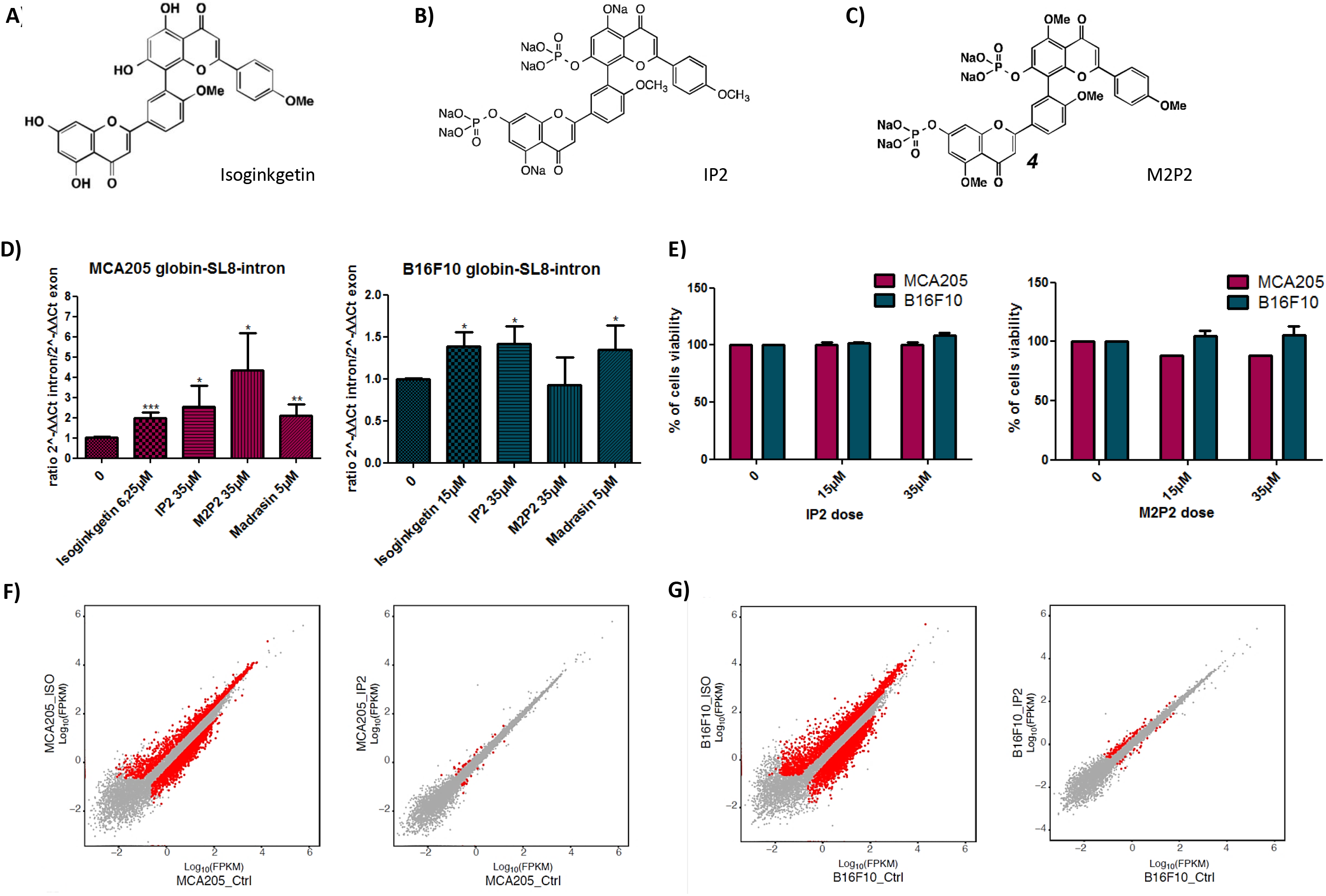
Synthesis and activity of the isoginkgetin derivatives IP2 and M2P2. Molecular structure of **(A)** isoginkgetin, **(B)** IP2 and **(C)** M2P2 compounds. **(D)** qRT-PCR analysis of splicing inhibition in mouse sarcoma MCA205 (left panel) and mouse melanoma B16F10 (right panel) cell lines both transiently expressing the globin-SL8-intron construct and treated with 15μM of isoginkgetin, 5μM of madrasin, 35μM of IP2 or 35μM of M2P2 for 48 hours. Data are given as mean ± SEM of the ratio of 2^-ΔΔCt^ intron to 2^-ΔΔCt^ exon of at least three independent experiments. *P<0.05, **P<0.01, ***P<0.001 (unpaired student t test). **(E)** Toxicity of IP2 (left panel) and M2P2 (right panel) compounds against MCA205 and B16F10 cells assessed by MTT assay. Data are expressed as mean ± SEM of the percentage of viable cells compared to the control condition of at least three independent experiments. **(F)** Comparison of gene expression profile between isoginkgetin-treated (left panel) and IP2-treated (right panel) MCA205 sarcoma cells. **(G)** Comparison of gene expression profile between isoginkgetin-treated (left panel) and IP2-treated (right panel) B16F10 melanoma cells. FPKM, Fragments per kilobase of exon per million reads mapped.

The water solubility of IP2 and M2P2 was found to be considerably higher than that of the parent compound isoginkgetin (data not shown). In addition, we tested the ability of IP2 and M2P2 to inhibit the splicing of the Globin-SL8-intron gene product in MCA205 and B16F10 cells. Interestingly, IP2 and M2P2 provide two distinct patterns of splicing inhibition in each cell line. IP2 treatment increases the presence of non-spliced RNA products in both cell lines, just as isoginkgetin or madrasin do. In contrast, M2P2 treatment does not impact splicing in B16F10 cells while it has a strong impact on splicing in MCA205 compared to IP2, isoginkgetin and madrasin treatment (Fig. 3D). Hence, IP2 and M2P2 seem to inhibit the splicing machinery in different ways. Importantly, we observed that the splicing pattern of IP2 is comparable to that of isoginkgetin and madrasin for the studied gene product. Finally, both IP2 and M2P2 compounds display no toxicity toward MCA205 and B16F10 WT cells at the doses tested as assessed by MTT assay (Fig. 3E). To further support the absence of toxicity of IP2, we performed from RNA-seq data a differential gene expression analysis on MCA205 and B16F10 cells treated with IP2 and compared it with isoginkgetin. We observed that IP2 impacts the expression of a highly reduced number of genes compared to the parental natural product isoginkgetin which shows cytotoxic effect (Figs. 3F and 3G). The greater selectivity of IP2 can explain the absence of toxicity on both cell lines. Overall, we have provided two new drugs that are water soluble and that impact the splicing differently in our two model cell lines at doses that do not impair cell viability.

### The isoginkgetin derivative IP2 efficiently increases MHC-I presentation of intron-derived antigen *in vitro*, reduces tumor growth *in vivo* and extends survival

IP2 and M2P2 compounds were first tested for their ability to increase the MHC-I presentation of PTP-derived antigens *in vitro*. For that purpose, MCA205 and B16F10 cells were transiently expressing the Globin-SL8-intron construct and treated with 15μM or 35μM of IP2 or M2P2. While treatment with IP2 increases the intron-derived SL8-antigen presentation in MCA205 and B16F10 cells similarly to what we observed upon isoginkgetin treatment (Figs. 4A and 4B, left panels), M2P2 decreases its presentation in MCA205 cells and does not impact it in B16F10 cells (Figs. 4A and 4B right panels). These results are interestingly correlated to the respective ability of IP2 and M2P2 to inhibit the splicing of the Globin-SL8-intron gene. In fact, M2P2 has no impact on both the splicing and the SL8 antigen presentation in B16F10. Conversely, M2P2 strongly inhibits the splicing in MCA205 and negatively affects the SL8 presentation. Hence, a tight regulation of splicing is required to positively impact the production and the presentation of intron-derived epitopes. Along with these results, we showed that the expression of H2-K^b^ molecules at the cell surface is not affected in the cell lines treated with IP2 and M2P2 (Supplementary Fig. S6A). In addition, looking at the cell surface K^b^ re-expression after acid stripping in IP2-treated MCA205 cells, we observed neither an increase in the kinetics of recovery of K^b^ molecules at the cell surface nor a higher level of expression at steady state (Supplementary Fig. S6B). This result supports the idea that the effect of IP2 on antigen presentation is earlier than the loading of the antigenic peptides onto the MHC class I molecules and the export of the peptide/MHC class I complex at the cancer cell surface. We can further conclude that the upregulation of PTP-dependent cancer immune response after IP2 treatment is not due to an overall increase of cell surface MHC class I K^b^ molecules. Finally, we confirmed that IP2 does not induce apoptosis of tumor cells even at a high dose (Supplementary Figs. S6C and S6D). These results show that the isoginkgetin derivative IP2 acts as a booster of the PTP-derived antigen production and presentation *in vitro* and *in vivo* in the same way as the natural product.

**Figure 4:**
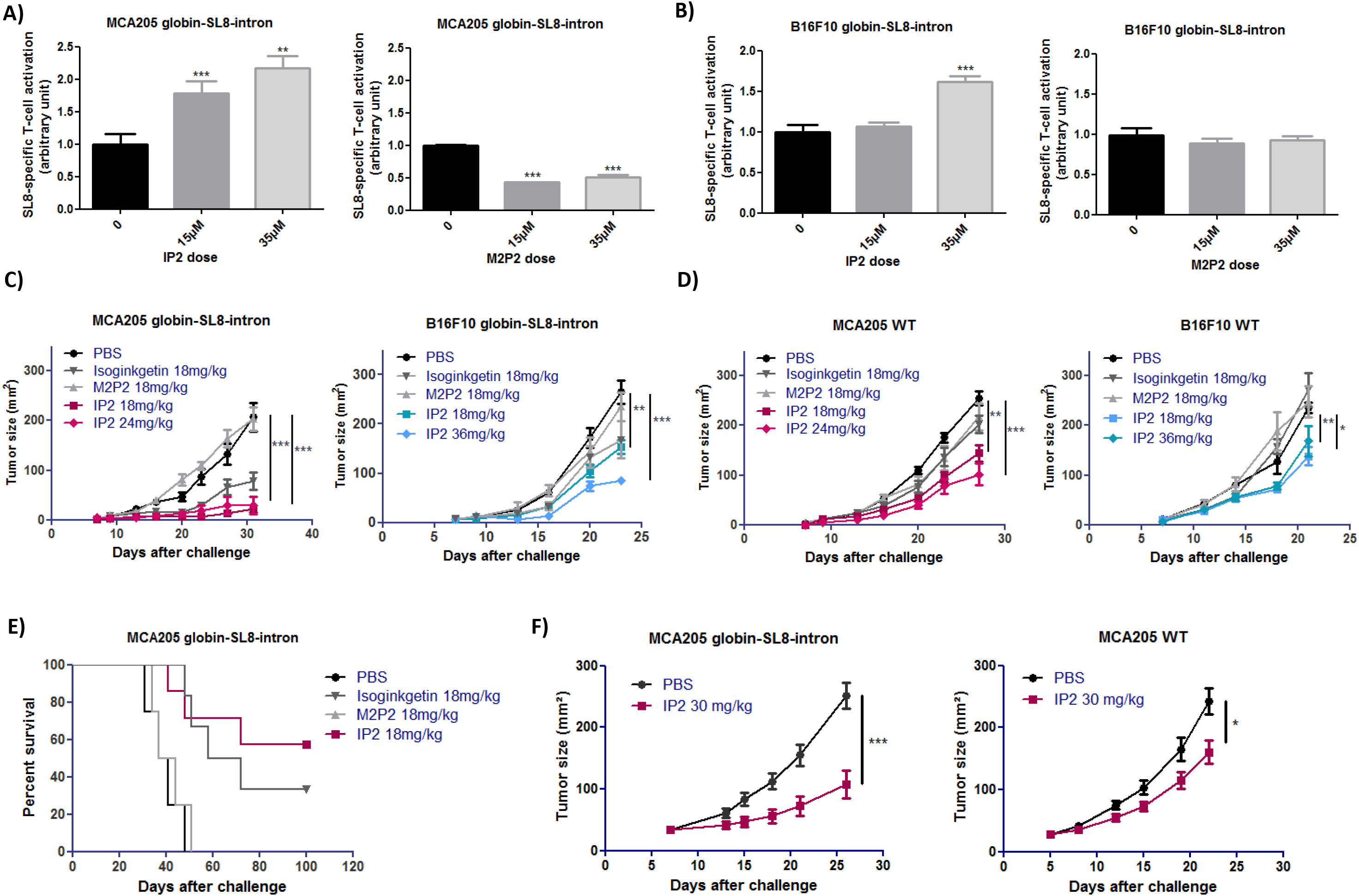
IP2 treatment reduces tumor growth and extends survival. B3Z SL8-specific T-cell activation after co-culture with **(A)** mouse sarcoma MCA205 and **(B)** mouse melanoma B16F10 cell lines both transiently expressing the intron-derived SL8 epitope and treated with 15μM or 35μM of IP2 (left panel) or M2P2 (right panel) for 18 hours. Free soluble SL8 peptide was added to ensure that T-cell activation assays were carried out in non-saturated conditions. Unspecific B3Z T-cell activation was taking into account in the results. Each graph is one representative of at least four independent experiments. Data are given as mean ± SEM. *P<0.05, **P<0.01, ***P<0.001 (unpaired student t test). Growth of **(C)** mouse sarcoma MCA205 (left panel) and mouse melanoma B16F10 (right panel) both stably expressing the globin-SL8-intron construct and **(D)** MCA205 WT (left panel) and B16F10 WT (right panel) cells subcutaneously inoculated into the right flank of immunocompetent C57BL/6 mice thereafter treated intraperitoneally with 18mg/kg of isoginkgetin, 18 mg/kg of M2P2, or with 18mg/kg, 24mg/kg or 36mg/kg of IP2 at day 5, 10 and 15 post tumor inoculation. Tumor size was assessed every 3 to 4 days until it reached the established ethical endpoints. Each line represents the tumor size in area (mm^2^) of at least 6 mice in each group. Data are given as mean ± SEM. *p<0.05, **p<0.01 (ANOVA with Tukey’s multiple comparison test comparing all groups). **(E)** Kaplan-Meier survival plot of mice subcutaneously inoculated with MCA205 sarcoma cells stably expressing the globin-SL8-intron construct and thereafter treated intraperitoneally with 18mg/kg of isoginkgetin, 18mg/kg of M2P2 or 18mg/kg of IP2. A Log-rank (Mantel-Cox) test was performed. **(F)** Growth of mouse sarcoma MCA205 stably expressing the globin-SL8-intron construct (left panel) or MCA205 WT (right panel) subcutaneously inoculated into the right flank of immunocompetent C57BL/6 mice thereafter treated intraperitoneally with 30mg/kg of IP2 every three days once tumor size had reached 25-30mm^2^. Tumor size was assessed every 3 to 4 days until it reached the established ethical endpoints. Each line represents the tumor size in area (mm^2^) of at least 6 mice in each group. Data are given as mean ± SEM. *P<0.05, **P<0.01, ***P<0.001 (unpaired student t test).

Next, IP2 and M2P2 were tested for their antitumor effect *in vivo*. MCA205 and B16F10 tumor cells, stably expressing the Globin-SL8-intron construct or WT, were subcutaneously inoculated into mice as previously performed with isoginkgetin treatment. At day 5, 10 and 15 post tumor inoculation, each group of mice was intraperitoneally treated with 18mg/kg isoginkgetin, IP2 or M2P2. At this dose, a significant decrease in MCA205 globin-SL8-intron tumor growth was observed after treatment with IP2 comparable to the isoginkgetin treatment, while no impact of M2P2 treatment was monitored (Fig. 4C, left panel and Supplementary Fig. S7A, left panel). In addition, the reduction of B16F10 globin-SL8-intron tumor growth was similar after treatment with 18mg/kg of isoginkgetin or IP2, while M2P2 had no effect on tumor growth (Fig. 4C, right panel). As IP2 treatment reduces tumor growth and is highly soluble in water, we tried to increase the dose injected into mice in order to achieve a greater reduction of tumor growth. However, increasing the dose of IP2 did not improve the antitumor effect on MCA205 Globin-SL8-intron at 24mg/kg but it did enhance it on B16F10 Globin-SL8-intron at 36mg/kg (Fig. 4C, right panel and Supplementary Fig. S7A, right panel). Strikingly, isoginkgetin and M2P2 treatments did not impact the growth of either MCA205 WT or B16F10 WT tumors while IP2 treatment slows down both (Fig. 4D and Supplementary Fig. S7B). We observed a 40 to 60% reduction of the growth of MCA205 WT (Fig. 4D and Supplementary Fig. S6B, left panels) and a 40% reduction of the growth of B16F10 WT (Fig. 4D and Supplementary Fig. S6B, right panels). Moreover, IP2 treatment was shown to extend the survival of mice with more than 50% survivors 100 days after tumor inoculation (Fig. 4E). At the same time, around 30% of mice treated with isoginkgetin were still alive while M2P2 treatment did not improved survival. Finally, we evaluated the ability of IP2 to treat established tumors. Mice were inoculated with either 2×10^5^ MCA205 Globin-SL8-intron or 2×10^5^ MCA205 WT tumor cells. Once tumor size had reached 25- 30mm^2^ (around day 5 to 7 post tumor inoculation) mice were treated intraperitoneally with 30 mg/kg of IP2 every three days until a total of 4 injections. At this dose, in mice bearing SL8-expressing MCA205 tumors, we observed a significant reduction of tumor size, over 65% at day 27 after challenge (Fig. 4F, left panel) while in mice bearing MCA205 WT tumors, we observed a significant reduction of tumor size, over 25% at day 27 after challenge (Fig. 4F, right panel).

Overall, these results suggest a correlation between the increase in PTP-derived antigen presentation observed *in vitro* and the reduction of the tumor growth *in vivo* upon treatment. Interestingly, contrary to isoginkgetin, IP2 treatment slows down the growth of tumors that do not bear the highly immunodominant SL8 epitope derived from PTPs. The difference of efficacy between the two molecules can be due to their biodisponibility, which should be higher for the water-soluble IP2 than for the hydrophobic isoginkgetin, as well as to the use of an increased dose of treatment. We suggest that the splicing inhibitor IP2 potentiates the cell surface apparition of immunodominant epitopes, driving the antitumor response.

### IP2 action is dependent on the immune response and creates a long-lasting antitumor response

To determine the requirement of the immune system and especially of the T-cell response for IP2 efficacy against tumors, we looked at its effect in Nu/Nu athymic nude mice that lack T-cells but not B and NK cells. As previously tested with isoginkgetin, MCA205 or B16F10 tumor cells stably expressing the Globin-SL8-intron construct or WT were subcutaneously inoculated into mice. At day 5, 10 and 15 post tumor inoculation, each group of mice was intraperitoneally treated with the dose of IP2 which was found to be the most efficient in immunocompetent mice. Hence, mice bearing MCA205 Globin-SL8-intron, MCA205 WT and B16F10 Globin-SL8-intron or WT tumors were treated with 18, 24 and 36 mg/kg respectively. In each condition, no impact of IP2 treatment was observed on tumor growth (Figs. 5A and 5B).

In addition, we tested the impact of *in vivo* CD8+ T cell depletion. Mice were subcutaneously inoculated with MCA205 Globin-SL8-intron or WT cells and thereafter treated with anti-CD8+ T cell antibody or with the isotype control. IP2 treatment was administered as previously at day 5, 10 and 15 post tumor inoculation. Interestingly, CD8+ T cell depletion completely abrogated the antitumor effect of the IP2 treatment (Fig. 5C). Therefore, this result confirms that the effect of the IP2 treatment on tumor growth is dependent on the CD8+ T cell response, which supports an antigen-driven cytotoxic activity against the tumor cells.

**Figure 5:**
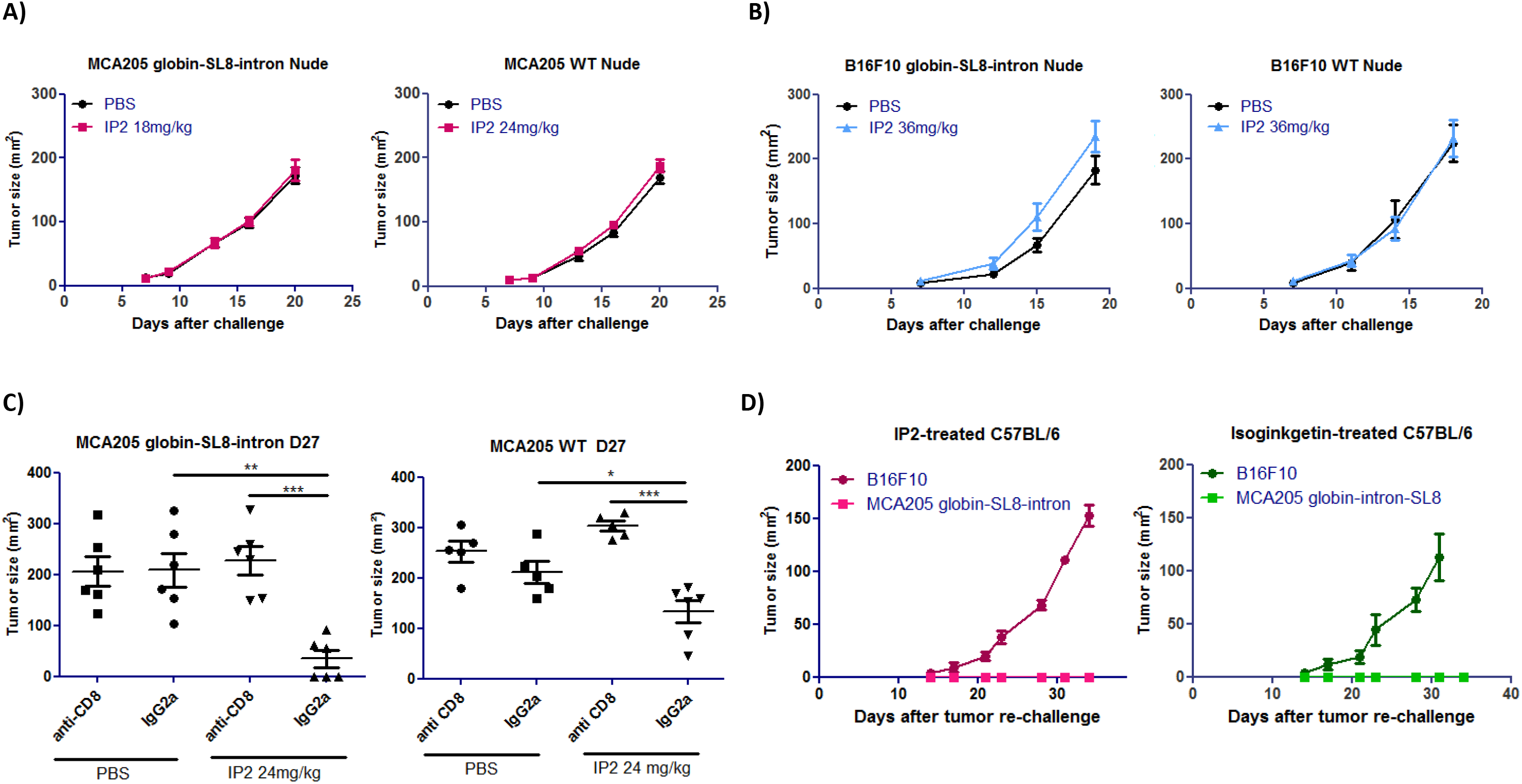
IP2 therapeutic effect is dependent on the immune response. Growth of **(A)** mouse sarcoma MCA205 stably expressing the globin-SL8-intron construct (left panel) or MCA205 WT (right panel) and **(B)** mouse melanoma B16F10 stably expressing the globin-SL8-intron construct (left panel) or B16F10 WT (right panel) subcutaneously inoculated into the right flank of immunodeficient Nu/Nu mice thereafter treated intraperitoneally with 18mg/kg of IP2 at day 5, 10 and 15 post tumor inoculation. Tumor size was assessed every 3 to 4 days until it reached the established ethical endpoints. Each line represents the tumor size in area (mm^2^) of at least 6 mice in each group. Data are given as mean ± SEM. *p<0.05, **p<0.01 (ANOVA with Tukey’s multiple comparison test comparing all groups). **(C)** Tumor size in area of mouse sarcoma MCA205 stably expressing the globin-SL8-intron construct (left panel) and MCA205 WT (right panel) subcutaneously inoculated into the right flank of immunocompetent C57BL/6 mice thereafter treated with 24mg/kg of IP2 at day 5, 10 and 15 post tumor inoculation and anti-CD8+ T cell depleting antibody every three days. Data are given as mean ± SEM. *p<0.05, **p<0.01 (ANOVA with Tukey’s multiple comparison test comparing all groups). **(D)** Growth of MCA205 cells stably expressing the globin-SL8-intron construct and B16F10 WT cells inoculated at day 100 into the right and left flanks respectively of C57BL/6 mice which experienced total tumor regression after first inoculation of MCA205 globin-SL8-intron at day 0 and treatment with IP2 (left panel) or isoginkgetin (right panel). Each line represents the tumor size in area (mm^2^) of at least 4 mice in each group.

Finally, around 50% of mice inoculated with MCA205 Globin-SL8-intron and subsequently treated with IP2 as described above experienced complete tumor rejection. 100 days after the first tumor inoculation, these mice were re-challenged with the same MCA205 Globin-SL8-intron tumor cells on the right flank and with B16F10 WT cells on the left flank. While B16F10 WT tumor cells grew over time, MCA205 Globin-SL8-intron tumor cells did not grow in mice. These results demonstrate that mice developed a long term antitumor response against SL8-expressing tumors after IP2 treatment.

Overall these results shed light on the capacity of specific splicing inhibitors, such as isoginkgetin and IP2, to positively modulate the antitumor immune response. In addition, they confirm that PTP-derived antigens are efficiently presented and recognized by CD8+T cells *in vitro* and *in vivo* and that a change in their presentation at the cell surface in quantity or in quality can lead to a CD8+ T cell-mediated response against cancer.

### IP2 treatment alters the MHC I immunopeptidome of tumor cells and modifies the presentation of both conventional and non-conventional MIPs

To confirm that the enhanced immune response observed after IP2 treatment results from a change in cancer cell immunogenicity, we analyzed the peptide repertoire displayed on the cancer cell surface upon treatment. MCA205 and B16F10 cells were cultured with or without IP2 and peptides eluted from H-2K^b^ and H-2D^b^ molecules were sequenced by mass spectrometry. We first observed that in the presence of IP2 the MHC I immunopeptidome of both MCA205 and B16F10 cells was altered. We identified peptides significantly enriched on the cell surface of MCA205 and B16F10 cells upon treatment and others whose presence was reduced (Fig. 6). Interestingly, we found peptides which have been already described in the immune epitope database IEDB but also peptides derived from retained introns (Table 1). This result confirms that retained introns can be a source of peptides for the MHC I presentation pathway and that allegedly non-coding regions of the genome can play a role in tumor cell immunogenicity. Furthermore, table 2 lists the peptides found exclusively after IP2 treatment in the three replicates of the peptide elution experiment. Here again, we found both exon-derived and retained intron-derived peptides, most of which have never been described before. We then seek to further characterize the peptides found exclusively after IP2 treatment. In B16F10 cells, there was a significant change in the length of peptides with 8-mers and 9-mers being enriched at the expense of both 11-mers and 12-mers (Supplementary Fig. S8). We then computed the predicted MHC I binding affinity of the peptides found exclusively on IP2-treated cells using the NetMHC 4.0 server. Peptides displayed on the surface of IP2-treated B16F10 cells are stronger binders of MHC-I H-2K^b^ and MHC-I H-2D^b^ molecules (Supplementary Fig. S8). There were twice more peptides with IC_50_ less than 500nM in IP2-treated cells compared to control ones. The same trend was observed in MCA205 cells but did not reach significance. Taken together those results show that IP2 re shapes the MHC-I immunopeptidome of cancer cells and induces the presentation of a new repertoire of epitopes, from both coding and allegedly non-coding sequences, which may promote tumor cell recognition by the immune system and tumor eradication.

**Figure 6:**
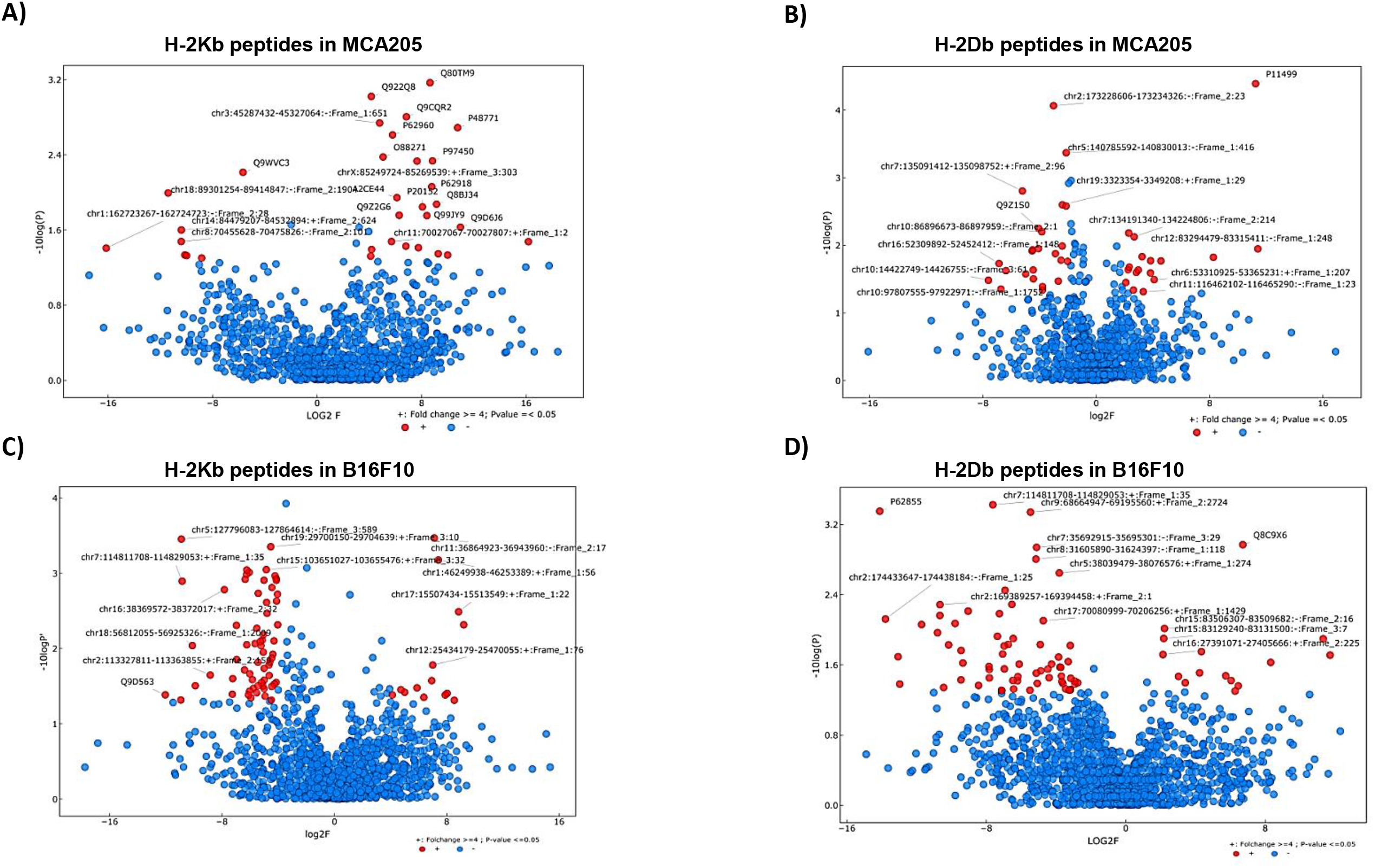
IP2 alters the MHC-I immunopeptidome of tumor cells. Volcano plots displaying the differential presentation of **(A)** H-2Kb peptides in MCA205 cells, **(B)** H-2Db peptides in MCA205 cells, **(C)** H-2Kb peptides in B16F10 cells and **(D)** H-2Db peptides in B16F10 cells between IP2-treated and untreated cells. The vertical axis corresponds to the mean expression value of logı_0_ (p-value) and the horizontal axis displays the log 2 fold change value. Red dots represent the peptides displaying at least a four-fold change with a p-value less than 0.05 between the untreated vs IP2-treated triplicates. Positive x-values represent up-regulation and negative x-values represent down-regulation.

**Table 1:**
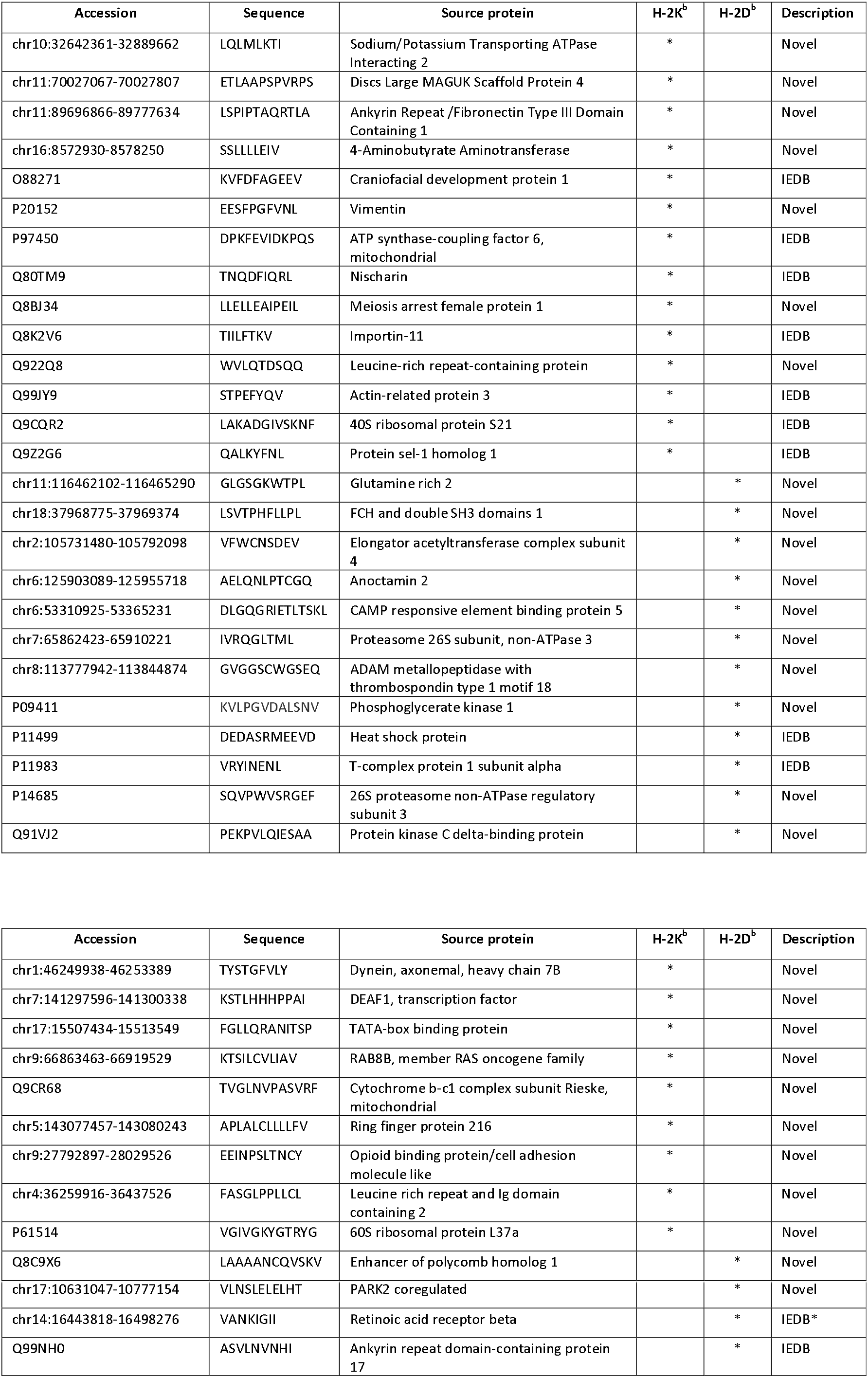
The MHC-I immunopeptidome of IP2-treated cells is enriched in peptides derived from both exons and retained introns. List of peptides displaying at least a four-fold increase with a p-value less than 0.05 between the untreated vs IP2-treated triplicates (red dots in the right half of the volcano plots in fig. 6) in MCA205 cells (upper table) and B16F10 cells (lower table). Accession numbers refer either to the *Mus Musculus* UniProt or to our in house *Retained Introns* databases (cf Material and methods). “Novel” indicates that the present study is the first time the peptide has been reported. “IEDB” indicates that the peptide is found in the immune epitope database. “IEDB*” indicates that the peptide is found in the immune epitope database associated with a different source protein. Peptides highlighted in grey are referred as “cancer-related genes” in the Protein Atlas.

**Table 2:**
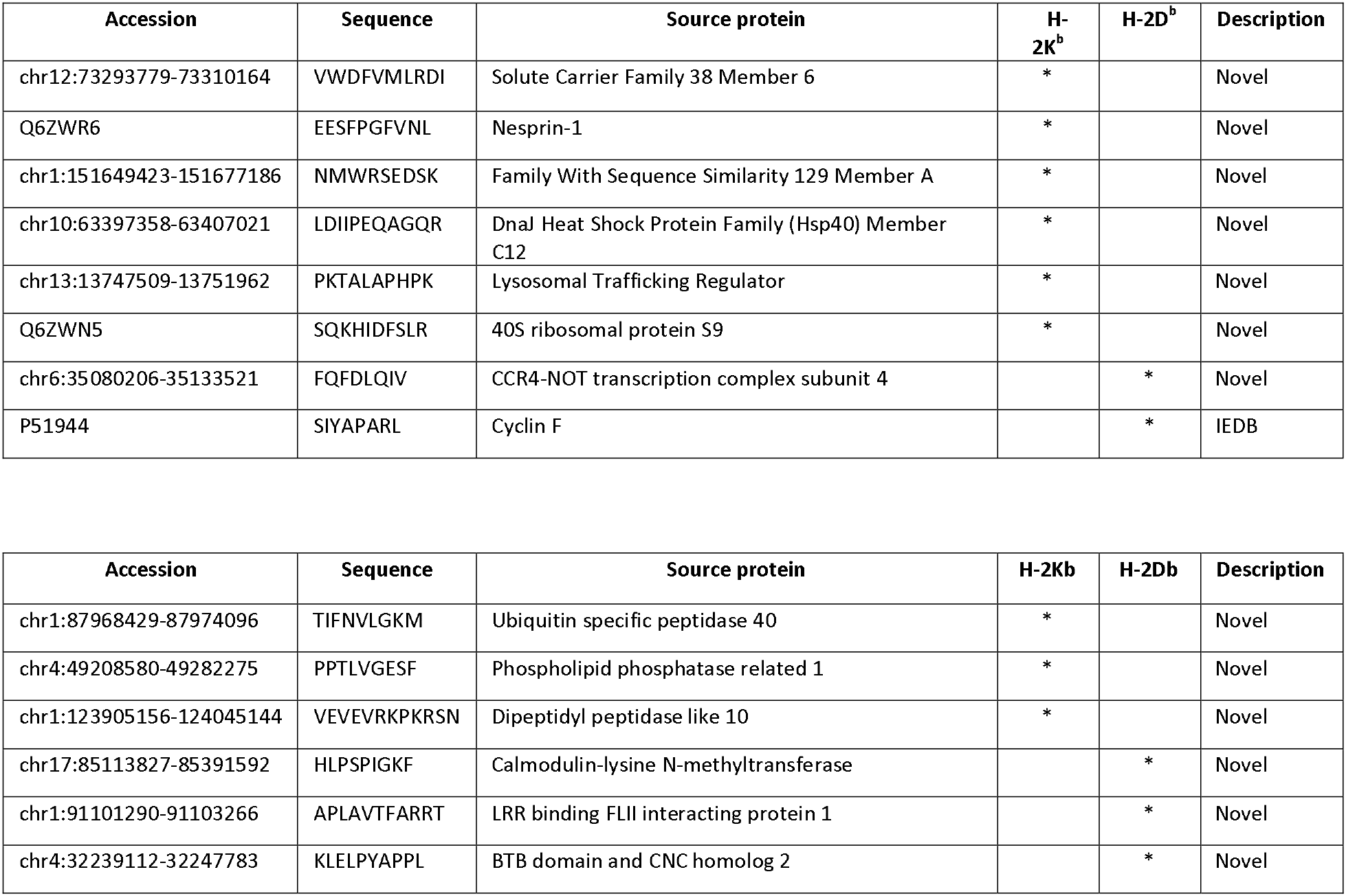
IP2 treatment leads to the presentation of new peptides derived from both exons and retained introns. List of peptides found exclusively after IP2 treatment in MCA205 cells (upper table) or B16F10 cells (lower table) with a p-value less than 0.05 between the untreated vs IP2-treated triplicates. Accession numbers refer either to the *Mus Musculus* UniProt or to our in house *Retained Introns* databases (cf Material and methods). “Novel” indicates that the present study is the first time the peptide has been reported. “IEDB” indicates that the peptide is found in the immune epitope database. Peptides highlighted in grey are referred as “cancer-related genes” in the Protein Atlas.

## Discussion

Splicing abnormalities have emerged as a specific feature of cancer cells and are studied as predictive markers for patient survival (30,31) as well as targets for cancer treatments with splicing inhibitors^23^, some of which are currently under development in acute myeloid leukemia (28). Here, we highlight splicing inhibitors as modulators of the adaptive antitumor immune response and suggest their action on PTP production and immunopeptidome modulation. We observed that treatment with the splicing inhibitor isoginkgetin or its derivative IP2 reduces the growth of sarcoma MCA205 and melanoma B16F10 tumors bearing the PTP-derived SL8 epitope and significantly extends mice survival. By performing the same experiment in Nude nu/nu mice, which are deficient in T cells but not in B or innate immune cells, we did not show tumor growth diminution upon treatment. Furthermore, *in vivo* depletion of CD8+ T cells in immunocompetent mice abrogates the tumor regression observed after IP2 treatment. We also observed that mice which experienced complete tumor rejection upon treatment with IP2 elicit a long lasting immune response. And finally, IP2 treatment was shown to reshape the MHC-I immunopeptidome, leading to a new antigenic peptide representation at the cell surface. Therefore, we conclude that an efficient adaptive immune response, particularly a CD8+ T cell-mediated response, is required for splicing inhibitors to impact tumor growth.

Because IP2 treatment has no impact on the growth of tumors in nude mice (Figure 5A) and does not induce apoptosis at the doses tested (Figure S6C and S6D), it is unlikely that IP2 induces a significant immunogenic cell death. The fact that isoginkgetin or IP2 impact on tumor growth is significantly higher in tumors expressing the PTP-derived SL8 epitope than the WT tumors demonstrates the key role of epitope immunogenicity. In addition, *in vitro* experiments provide evidence for these splicing inhibitors to induce a quantitatively higher number of presented SL8 peptides, driving the activation of specific T-cells. Conversely, the isoginkgetin derivative M2P2 does not increase the presentation of the SL8-derived PTPs *in vitro* and, unlike IP2 or isoginkgetin, does not impact the tumor growth *in vivo*, suggesting that both events may be correlated. Hence, we can reasonably hypothesize that IP2 or isoginkgetin treatments enhance the amount of SL8 peptide at the cancer cell surface *in vivo*, resulting in the induction of a higher T cell cytotoxic activity against tumor cells followed by tumor growth delay or tumor eradication. Besides, several mechanisms have been described where the presence of highly expressed antigen tumor variants allows the elimination of antigen loss variants, an effect which could explain the action of splicing inhibitors on tumor growth by an increased load of antigens in tumors (29,32–34).

The PTP model describes the pre-spliced mRNA as the template for the production of PTPs by an alternative translational event occurring in the nucleus. In the study describing this alternative translation, we demonstrated previously that forced nuclear retention of the mRNA encoding the intronic SL8 peptide leads to an increase in the SL8 antigen presentation (18). Overall, splicing inhibition has been shown to promote pre-RNA accumulation in the nucleus (35,36). The link between nuclear accumulation of pre-mRNAs and increased antigen presentation is not known so far. It is tempting to hypothesize that accumulation of pre-mRNA in the nucleus provides more templates for the pioneer round of translation leading to the enrichment of SL8-containing PTPs used as a major source for SL8 direct presentation.

The fact that madrasin, isoginkgetin and IP2, but not M2P2, positively impact PTP-derived antigen presentation shows that a tight regulation of splicing is needed for this effect and encourages future investigations on the link between splicing and antigen presentation. In fact, while M2P2 impacts the splicing of the Globin-SL8-intron gene product in MCA205 cells in a greater extent than isoginkgetin, madrasin and IP2 molecules (Figure 3E), it does not increase MCA205 antigen presentation *in vitro* and does not impact tumor growth *in vivo* (Figure 4), adding more complexity to the correlation between splicing inhibition and antigen presentation. Since most of the splicing inhibitors appear to affect the formation of the basal splicing machinery, one might expect that such modulator affect globally total mRNA splicing. However, it has been reported that there is a specificity of action of each splicing inhibitor depending on certain groups of genes (37,38). Therefore, these differences of action between IP2 and M2P2 in terms of antigen presentation and inhibition of the spliceosome suggest that M2P2 might act through a different mechanism on a group of genes that is not affected by IP2 treatment. Furthermore, while isoginkgetin, madrasin and IP2 impacts similarly the splicing of the Globin-SL8-intron gene product in B16F10 cells, M2P2 does not impact the splicing in this cell line (Figure 3D), which proves that the splicing inhibitory effect is cancer specific and that each splicing inhibitor can have specific targets. Cancer cells can acquire deficiencies in the splicing machinery that benefit their growth, for example by preventing the expression of tumor suppressor genes^23^. These deficiencies do not have the same nature in all tumors and therefore could explain the distinct effects of splicing inhibitors on separate tumor types. These results emphasize the need for assessing the specificity of each splicing inhibitor on different cancer types and group of genes.

Cytotoxic T lymphocytes failure to reject tumors can in part be explained by an initial inappropriate activation of CTL by pAPCs. We recently demonstrated that tumor-associated PTPs are a source material for CD8+ T cell cross-priming by dendritic cells (DCs) and provided hints that PTPs for endogenous and cross-presentation are produced by the same translational event followed by the quick separation of the two pathways^17^. PTPs are rare products^25^ and the efficiency of PTP vaccines was shown to rely on the previous enrichment of PTPs with proteasome inhibitor^17^. Therefore, splicing inhibitors IP2 and isoginkgetin may, in addition to provide more source materials for the direct antigen presentation, enrich the pool of SL8-containing PTPs that serve as a source material for intratumoral DC uptake and cross-presentation, inducing an enhanced proliferation of SL8-specific CD8+ T cells. Intriguingly, IP2 treatment slows down the growth of WT tumors, suggesting that an immunodominant epitope is not required to observe an impact on tumor but however needed to observe complete tumor regression. While a quantitative change of naturally expressed epitope in WT tumors upon treatment may be involved in this effect, qualitative changes may also occur. The plasticity of the immunopeptidome has been highlighted in several studies (39) and has been linked to many parameters such as intracellular metabolism and transcriptional regulations, gene expression and genetic polymorphism (40–43). In fact, we demonstrate that interfering with the spliceosome using the IP2 compound leads to a reshaping of the MHC-I immunopeptidome of cancer cells (Figure 6). This IP2-induced immunopeptidome is composed not only of antigenic epitopes whose expression has been increased compared to the non-treated cells but also of new antigenic epitopes that are produced only upon treatment with IP2. In fact, Smart, Margolis and colleagues have recently suggested that retained introns can be a source of tumor-specific neoepitopes in cancer. By treating cells with IP2 we have also been able to demonstrate that the immunopeptidome of treated cancer cells is not only composed of epitopes coming from exonic sequences but also from retained introns. We therefore confirmed that retained intron can be an important source of tumor neoepitopes. Consequently, IP2 treatment is likely to induce the presentation of new MIPs recognized by CD8+ T cells, delaying the tumor growth. The presence of peptides derived from “cancer-related” proteins of the Protein Atlas, such as heat shock proteins, metallopeptidases and oncogenes, suggests that IP2 treatment could force the tumor cells to present peptides facilitating their recognition by the immune system. The apparition of those new MIPs might result from the translation of pre-spliced mRNAs enriched after IP2 treatment leading to the apparition of more MIPs derived from non-coding regions. The latter have been suggested to be frequently mutated in cancer (44) and may extend far beyond retained introns to 5 ‘ and 3 ‘UTR, spliced introns, long non-coding RNAs and many others.

While a deeper understanding of isoginkgetin and IP2 treatments on splicing is needed, their role during immune response opens a large field of study on the link between splicing and immunopeptidome that could lead to the development of innovative anti-cancer therapies targeting the spliceosome.

## Material and methods

### Cell culture, transfection, plasmids and drugs

MCA205 mouse sarcoma cell line is cultured at 37°C under 5% CO_2_ in RPMI 1640 medium (Life Technologies) in the presence of 1% glutamine, 1% sodium pyruvate, 1% non-essential amino-acids, 1% penicillin/streptomycin and 10% FBS (Life Technologies) under standard conditions. B16F10 mouse melanoma cell line, MRC5 human fibroblast cell line and A375 human melanoma cell line are cultured at 37°C under 5% CO_2_ in DMEM medium (Life Technologies) containing 1% glutamine, 1% penicillin/streptomycin and 10% FBS. A549 human lung carcinoma cell line is cultured under standard conditions in DMEM/F12 + Glutamax I in the presence of 1% Hepes, 1% sodium pyruvate and 10% FBS. Stable MCA205-Globin-SL8-intron and B16F10-Globin-SL8-intron cell lines are cultured under the same condition as MCA205 and B16F10 cell lines respectively, with additional 2mg/ml G418 (Life Technologies) for selection. Stable MCA205-Ovalbumin and B16F10-Ovalbumin cell lines are cultured under the same condition as MCA205 and B16F10 cell lines respectively, with additional 0.2μg/ml zeocin (Life Technologies) for selection. The SL8/H-2K^b^-specific (B3Z) T-cell reporter hybridoma (45) are cultured at 37°C under 5% CO2 in RPMI 1640 medium in the presence of 1% glutamine, 0.1% β-galactosidase, 1% penicillin/streptomycin and 10% FBS under standard conditions. MCA205, A375, A549 and MRC5 cell lines are transfected with the reagent jetPRIME (Ozyme) according to the manufacturer protocol. B16F10 cell line is transfected with the reagent GeneJuice (Millipore) according to the manufacturer protocol. The plasmids YFP-Globin-SL8-intron, YFP-Globin-SL8-exon and PCDNA3-Ova have been described previously^25^. The pre-mRNA splicing inhibitor isoginkgetin from Merck Millipore and the madrasin from Sigma-Aldrich are dissolved in DMSO according to the manufacturer protocol. IP2 and M2P2 were synthesized from the isoginkgetin powder and dissolved in PBS.

### T-cell activation assay

MCA205 and B16F10 mouse cell lines are transfected with the plasmid YFP-Globin-SL8-intron or with the PCDNA3 empty plasmid (negative control) with the transfection reagent jetPRIME (Ozyme) or GeneJuice (Millipore) respectively according to each manufacturer protocol. A375, A549 and MRC5 human cell lines are transfected with the plasmid encoding mouse H2-K^b^ molecule for 12 hours followed by the transfection of the plasmid YFP-Globin-SL8-intron with the transfection reagent jetPRIME (Ozyme) according to the manufacturer protocol. Twenty-four hours after transfection, cells are treated with different doses of Isoginkgetin (Merk Millipore), IP2 or M2P2 overnight. Then cells are washed three times with PBS 1X and 5×104 cells are co-cultured with 1×105 B3Z cells. In positive control wells, 4μg/ml of synthetic peptide SL8 is added. Cells are then incubated at 37°C with 5% CO2 overnight. Cells are centrifuged at 1200rpm for 5min, washed twice with PBS 1X and lysed for 5min at 4°C under shaking with the following lysis buffer: 0.2% TritonX-100, 10mM DTT, 90 mM K2HPO4, 8,5 mM KH2PO4. The lysate is centrifuged at 3000rpm for 10min and the supernatant is transferred to a 96-well optiplate (Packard Bioscience, Randburg, SA). The revelation buffer containing 33mM of methylumbellifery β-D-galactopyranoside (MUG) is added and the plate is incubated at room temperature for 3 hours. Finally, the β-galactosidase activity is measured using the FLUOstar OPTIMA (BMG LABTECH Gmbh, Offenburg, Germany). Results are expressed as mean ± SEM. *P<0.05, **P<0.01, ***P<0.001 (unpaired student t test).

### FACS Analysis: H-2Kb expression and recovery at the cell surface

Mouse cells are treated with drugs for 18hours. Human cells are transfected with the plasmid encoding the mouse H-2K^b^ molecule for 24 hours, followed by 18hours treatment with drugs. Cells are harvested and 2×10^6^ human cells are stimulated with 5μg SIINFEKL synthetic peptide for 15min at 37°C, 5% CO2. Human and murine cell lines are incubated with human or mouse FcR block respectively for 10min at 4°C according to the manufacturer protocol (Miltenyi Biotech). Mouse cells are stained with anti-mouse MHC class I (H-2K^b^)-PerCP-eFluor®710 (AF6-88.5.5.3, eBioscience) or with the corresponding mouse IgG2a K isotype control-PerCP eFluor®710 (eBM2a, eBioscience) for 30min at 4°C. Human cells are stained with anti-mouse OVA257-264 (SIINFEKL) peptide bound to H-2K^b^-APC (25-D1.16, eBioscience) for 30min at 4°C. Cells are stained with DAPI prior to acquisition for dead cell exclusion. In mouse cell lines the isotype antibody is used for PerCP eFluor®710-positive gating while in human cell lines non-stimulated cells are used for APC-positive gating. To study the kinetics of endogenous surface K^b^ recovery, cells were treated for 18h with the different drugs. The day after, cells were washed and treated with ice-cold citric acid buffer (0.13M citric acid, 0.061M Na_2_HPO_4_, 0.15M NaCl [pH 3]) at 1×10^7^ cells per milliliter for 120s, washed three times with PBS, and resuspended in culture medium. At the indicated time point, an aliquot of cells (generally 1.5×10^6^) was removed and stained with anti-mouse H-2K^b^ PE or anti-mouse SIINFEKL-peptide bound to H-2K^b^-PE. All flow cytometry experiments were conducted using the BD LSRII flow cytometer (BD Biosciences) and data are analyzed with the FlowJow software (V10). Results are expressed as mean ± SD. *P<0.05, **P<0.01, ***P<0.001 (unpaired student t test).

### Annexin V staining

Mouse cells are treated with drugs for 18hours. Floating and adherent cells are collected, washed in PBS and resuspended in 1X Annexin V Binding Buffer with APC Annexin V (BD Pharmingen) according to the manufacturer protocol. Cells are gently vortexed and incubated 15 min at RT in the dark. Cells are stained with DAPI prior to flow cytometry analysis. Cells are analyzed using the BD LSRII flow cytometer (BD Biosciences) and data are analyzed with the FlowJow software (V10). Results are expressed as mean ± SEM. *P<0.05, **P<0.01, ***P<0.001 (unpaired student t test).

### RNA preparation, RT and qPCR

Cells are transfected with the YFP-Globin-SL8-intron plasmid for 24 hours followed by treatment with drugs for 48 hours. Cells are then harvested and total cellular RNA is extracted and purified using the RNeasy Mini kit (Qiagen) according to the manufacturer protocol. The reverse transcription is carried out with 0.5 μg of RNA with the iScript cDNA synthesis kit (Bio-Rad) according to the manufacturer protocol. The StepOne real-time PCR system (Applied BioSystems) is used for qPCR and the reaction is performed with the Power SYBR green PCR master mix (Applied BioSystems). The results are analyzed using the StepOne software. Mouse gene specific primers are designed as follow: 18S mouse forward primer, 5’-GCCGCTAGAGGTGAAATTCTTG-3’; 18S mouse reverse primer, 5’-CATTCTTGGCAAATGCTTTCG-3’; Glob-SL8-exon forward primer, 5’-AGAAGTCTGCCGTTACTGCC-3’; Glob-SL8-exon reverse primer, 5’-AGGCC AT C ACT AA AGGC ACC-3 ‘; Glob-SL8-intron forward primer, 5’-GTATCAAGGTTACAAGACAG-3’; Glob-SL8-intron reverse primer, 5’-GGGAAAATAGACCAATAGGC-3’.

### MTT assay

Cells are plated in a 96-wells plate with 1×10^4^ cells per well for 24 hours. Cells are then treated with drug overnight. After removing the medium from the plate, the MTT (3-[4,5-Dimethylthiazol-2-yl]-2,5-diphenyltetrazolium bromide) (Sigma-Aldrich) powder is re-suspended into PBS, filtered and added into the plate wells at 2,5mg/ml. The plate is incubated for 2 hours at 37°C, with 5% CO2. It is then centrifuged and the supernatant is carefully removed in order not to disturb the precipitated formazan. 200μl/well of DMSO is added and the plate is shaken for 10 minutes. Absorbance is read at 544nm using the FLUOstar OPTIMA. Each experiment is repeated three times and the average of the three results is expressed in percentage of cell viability compare to the untreated condition ± SEM.

### Animal studies: Mice tumor challenge and drug treatment

C57Bl/6J female mice are obtained from Harlan Laboratories. NU/NU nude mice are obtained from Charles River. 7 week-old mice are injected subcutaneously on the right flank with 5×10^4^ MCA205 wild type (WT) or 7.5×10^4^ MCA205-Globin-SL8-intron cells, or with 5×10^4^ B16F10 wild type (WT) or B16F10-Globin-SL8-intron cells along with matrigel (VWR). Five days after challenge, mice are treated intraperitonealy with different dose of PBS, Isoginkgetin, IP2 or M2P2. Ten and fifteen days after challenge mouse are again treated intraperitonealy with the same drug. Area of the tumor is recorded every 3 to 4 days until ethical limit points are reached. All animal experiments were carried out in compliance with French and European laws and regulations. Mice which experience complete tumor regression are kept for 100 days after tumor challenge and are then injected subcutaneously on the right flank with 7.5×10^4^ MCA205-Globin-SL8-intron cells and on the left flank with 5×10^4^ B16F10 WT cells. Results are expressed as mean ± SEM. *p<0.05, **P<0.01, ***P<0.001 (ANOVA with Tukey’s multiple comparison test comparing all groups).

### RNA-Seq Read Mapping and differential expression

MCA205 and B16F10 cells were treated with 15 μM of isoginkgetin or 35 μM of IP2 for 18h. RNA was extracted using QIAGEN RNeasy columns. RNA-seq libraries were prepared from total RNA by poly(A) selection with oligo(dT) beads according to the manufacturer’s instructions (Life Technologies). The resulting RNA samples were then used as input for first-strand stranded library preparation using a custom protocol based on dUTP method as described previously (1). Libraries were sequenced on the Illumina HiSeq 4000 sequencer using 50-base pair paired-end read method.

All RNA-seq data were aligned to the mouse genome assembly mm10 with the TopHat2 splice-junction mapper for RNA-Seq reads (version 2.1.1) with default parameters and the Cuffdiff program (version 2.2.1) for all analyses of differential expression. CuffDiff was run using the following parameters: –library-type=fr-firststrand –compatible-hits-norm –library-norm-method geometric – min-reps-for-js-test 2 –dispersion-method per-condition -u -b. In all such analyses, the difference in expression of a gene was considered significant if the q value was less than 0.05 (the Cuffdiff default) and the fold change was superior to 2. Scatter plots were generated using ggplot2 in R environment.

### RNA-Seq and generation of the *Retained Introns* database

Libraries were prepared with TruSeq Stranded Total RNASample preparation kit according supplier recommendations. Briefly the key stages of this protocol are successively, the removal of ribosomal RNA fraction from 1μg of total RNA using the Ribo-Zero Gold Kit, a fragmentation using divalent cations under elevated temperature to obtain approximately 300bp pieces, double strand cDNA synthesis, using reverse transcriptase and random primers, and finally Illumina adapters ligation and cDNA library amplification by PCR for sequencing. Sequencing is then carried out on paired-end 75b of Illumina HiSeq4000. The *Retained Introns* database was generated from RNA-seq data (4 replicates for each cell line) according to KMA (KeepMeAround) pipeline which is a suite Python scripts and a R package. Contrary to other existing tools, KMA improves its accuracy by using biological replicates to reduce the number of false positive. The databases were built as follows: (i) Pre-processing step : Intron coordinates were generated by KMA scripts from the genome reference (GENCODE release GRCm38) and the associated annotation file (GTF file); (ii) Quantification step : RNA-seq files were mapped according to previously defined introns and transcript regions with Bowtie2 v2.3.3.1 and were then quantified with eXpress tool v1.5.2; (iii) Post-processing step : Previous generated files were processed by KMA to define a list of retained intron. All expressed introns were considered for the next step. Intron nucleic sequences are then translated in the first three reading frames to create the two databases. Generated amino-acid sequences are then separated according to stop codon and sequences with length lower than 8 amino-acid are discarded. Finally, we identified 135 348 retained introns for MCA205 cells and 140 943 retained introns for B16F10 cells where 124 911 were in common.

### HLA ligand elution

MCA205 and B16F10 cells were lysed in buffer containing PBS, 0.25% Sodium Deoxycholate and protease inhibitor (Pierce™ Protease Inhibitor Mini Tablets, EDTA-free, Thermo Fisher) during 1h at 4°C. Lysate was sonicated and centrifugation was performed 1h at 4500rpm at 4°C to pellet debris. Supernatant was filtered through 0.40 μm filter and applied on affinity columns overnight. Columns were prepared by coupling Y-3 and B22-249 antibodies to CNBr-activated Sepharose (GE Healthcare) (1 mg antibody/40 mg Sepharose). On the second day, the columns were eluted in 4 steps using 0.2% TFA. The eluate was filtered through a 3 kDa filter (Pall Nanosep^®^ centrifugal device with Omega membrane) and concentrated using a vacuum centrifuge.

### LC-MS/MS acquisition and qualitative analysis

The peptides were desalted using ZipTip C^18^ pipette tips (Pierce Thermo Scientific), eluted in 40 μL acetonitrile 70% (Fisher Scientific)/0.1% formic acid, vacuum centrifuged and resuspended in 12 μL of 0.1% formic acid. All C18 zip-tipped peptides extracts were analyzed using an Orbitrap Fusion Tribrid equipped with an EASY-Spray Nano electrospray ion source and coupled to an Easy Nano-LC Proxeon 1200 system (all devices are from Thermo Fisher Scientific, San Jose, CA). Chromatographic separation of peptides was performed with the following parameters: Acclaim PepMap100 C18 precolumn (2 cm, 75 μm i.d., 3 μm, 100 Å), Pepmap-RSLC Proxeon C18 column (75 cm, 75 μm i.d., 2 μm, 100 Å), 300nl/min flow, gradient rising from 95 % solvent A (water, 0.1% formic acid) to 28% solvent B (80% acetonitrile, 0.1% formic acid) in 105 minutes and then up to 40%B in 15min followed by column regeneration for 50 min. Peptides were analyzed in the orbitrap in full ion scan mode at a resolution of 120000 and with a mass range of *m/z* 400-650 using quadrupole isolation and an AGC target of 1.5×10^5^. Fragments were obtained by Collision Induced Dissociation (CID) activation with a collisional energy of 35 %. MS/MS data were acquired in the Orbitrap in a top-speed mode, with a total cycle of 3 seconds with an AGC target of 7×10^4^. The maximum ion accumulation times were set to 100 ms for MS acquisition and 150 ms for MS/MS acquisition in parallelization mode. All MS/MS data were processed with an in-house Sequest server node using Proteome Discoverer 2.2 (Thermo Scientific) for qualitative analysis. The mass tolerance was set to 10 ppm for precursor ions and 0.02 Da for fragments. The following modification was used as variable modification: oxidation. No enzyme search was applied. All peptides were filtered at 5% FDR. MS/MS data were searched against an in house *Retained Intron* database (MCA205 or B16F10) and Mouse Uniprot extracted database.

### Quantitative analysis in label-free experiments

Label-free quantification in between subject analysis was performed on raw data with Progenesis-Qi software 4.1 (Nonlinear Dynamics Ltd, Newcastle, U.K.) using the following procedure: (i) chromatograms alignment, (ii) peptide abundances normalization, (iii) statistical analyses of features, and (iv) peptides identification using Sequest server through Proteome Discoverer 2.2 (Thermo Scientific). The mass tolerance was set to 10 ppm for precursor ions and 0.02 Da for fragments. The following modification was used as variable modification: oxidation. No enzyme search was applied. A decoy search was performed and the significance threshold was fixed to 0.05. MS/MS data were searched against an in house *Retained Intron* database (MCA205 or B16F10) and Mouse Uniprot extracted database. The resulting files were imported into Progenesis-LC software.

## Data availability

The complete proteomics data sets are available in the PRIDE partner repository under the identification number: PXD012102 as.raw files, Proteome Discoverer 2.2.pdResult file, associated pep.xml and xlsx files, label-free peptides and proteins report generated by Progenesis QI.

## Acknowledgments

The A375 and A549 cell lines were a gift from Dr. Fabrice André, Gustave Roussy Institute. The MRC5 cell line was a gift from Dr. Patricia Kannouch, Gustave Roussy Institute. We also thank Prof. Laurence Zitvogel for valuable comments on the manuscript. This work was funded by The Avenir program (INSERM), La Fondation ARC, and Fondation Gustave Roussy. AP and RD are supported by the Université Paris Sud, MB is supported by the Fondation Gustave Roussy,

---

**Supplementary Fig. S1: Cell viability after treatment with isoginkgetin or madrasin**.

Viability of **(A)** human melanoma A375, human lung carcinoma A549 or normal human lung fibroblast MRC5 cell lines and **(B)** mouse sarcoma MCA205 and mouse melanoma B16F10 cell lines after treatment with isoginkgetin for 18h. Viability of **(C)** human melanoma A375, human lung carcinoma A549 or normal human lung fibroblast MRC5 cell lines and **(D)** mouse sarcoma MCA205 and mouse melanoma B16F10 cell lines after treatment with madrasin for 18h. Data are given as the mean of the percentage of cell viability ± SEM from at least three independent experiments. *P<0.05, **P<0.01, ***P<0.001 (unpaired student t test).

**Supplementary Fig. S2: Impact of cyclophosphamide and gemcitabine treatment on exon- and intron-derived antigen presentation in cancer cells.**

B3Z SL8-specific T-cell activation after co-culture with **(A)** human melanoma A375, human lung carcinoma A549 or normal human lung fibroblast MRC5 cell lines or **(B)** mouse sarcoma MCA205 and mouse melanoma B16F10 cell lines, all transiently expressing the mouse H2-K^b^ molecules and the intron-derived SL8 epitope and treated with 7μM or 21μM of cyclophosphamide for 18 hours. B3Z SL8-specific T-cell activation after co-culture with **(C)** human melanoma A375, human lung carcinoma A549 or normal human lung fibroblast MRC5 cell lines or **(D)** mouse sarcoma MCA205 and mouse melanoma B16F10 cell lines, all transiently expressing the mouse H2-K^b^ molecules and the intron-derived SL8 epitope and treated with 2.5μM or 6.25μM of gemcitabine for 18 hours. Free soluble SL8 peptide was added to ensure that T-cell activation assays were carried out in non-saturated conditions. Unspecific B3Z T-cell activation was taking into account in the results. Each graph is one representative of at least four independent experiments. Data are given as mean ± SEM. *P<0.05, **P<0.01, ***P<0.001 (unpaired student t test).

**Supplementary Fig. S3: Expression of H2-K^b^ molecules at the cell surface.**

**(A-D)** Gating strategies. Flow cytometry analyses of H2-K^b^ expression on mouse MCA205 and mouse B16F10 cells treated with **(B)** isoginkgetin or madrasin and **(C)** gemcitabine or cyclophosphamide. Flow cytometry analyses of H2-K^b^ expression on human melanoma A375, human lung carcinoma A549 or normal human lung fibroblast MRC5 cell lines, all transiently expressing the mouse H2-K^b^ molecules and treated with **(E)** isoginkgetin or madrasin and **(F)** gemcitabine or cyclophosphamide.

**Supplementary Fig. S4: Additional data from figure 2.**

Tumor size in area (mm2) of **(A)** mouse sarcoma MCA205 and **(B)** mouse melanoma B16F10 both stably expressing the globin-SL8-intron construct or **(C)** MCA205 WT and **(D)** B16F10 WT cells subcutaneously inoculated into the right flank of immunocompetent C57BL/6 mice thereafter treated intraperitoneally with 12mg/kg or 18mg/kg of isoginkgetin at day 5, 10 and 15 post tumor inoculation. Tumor size in area (mm2) of **(E)** mouse sarcoma MCA205 and **(F)** mouse melanoma B16F10 both stably expressing the globin-SL8-intron construct or **(G)** MCA205 WT and **(H)** B16F10 WT cells subcutaneously inoculated into the right flank of immunodeficient Nu/Nu mice thereafter treated intraperitoneally with 18mg/kg of isoginkgetin at day 5, 10 and 15 post tumor inoculation. Tumor size was assessed every 3 to 4 days until it reached the established ethical endpoints. Data are given as mean ± SEM. *p<0.05, **p<0.01 (ANOVA with Tukey’s multiple comparison test comparing all groups).

**Supplementary Fig. S5: Synthesis process of IP2 and M2P2 synthesis.** IP2 compound is referred as compound *2*. M2P2 compound is referred as compound *4*.

**Supplementary Fig. S6: IP2 does not impact H2-K^b^ molecules expression at the cell surface and does not induce apoptosis.**

**(A)** Flow cytometry analyses of H2-K^b^ expression on MCA205 (left panel) and B16F10 (right panel) cells treated with IP2 and M2P2. **(B)** Kinetics of the recovery of H-2Kb molecules at the cell surface of MCA205 cells after acid strip followed by flow cytometry with anti-H-2Kb antibody (left panel) or anti-SIINFEKL bound to H-2Kb antibody (right panel). Cells are treated with 25 μM of isoginkgetin, 35 μM of M2P2 or 35 μM of IP2 for 18h prior to acid strip. MFI, Mean Fluorescent Unit. **(C-D)** Apoptosis status of MCA205 and B16F10 cells treated with 35μM or 1000μM of IP2 for 18 hours assessed by DAPI/Annexin V staining.

**Supplementary Fig. S7: Additional data from Figure 4.**

Tumor size in area (mm^2^) of **(A)** MCA205 (left panel) and B16F10 (right panel) cells both stably expressing the globin-SL8-intron construct and **(B)** MCA205 WT (left panel) and B16F10 WT (right panel) cells subcutaneously inoculated into the flank of immunocompetent C57BL/6 mice thereafter injected intraperitoneally with 18mg/kg of isoginkgetin, 18mg/kg of M2P2 and 12mg/kg, 18 mg/kg, 24mg/kg or 36mg/kg of IP2 at day 5, 10 and 15 post tumor inoculation. Data are presented at the day when the ethical endpoints are reached and are given as mean ± SEM. *p<0.05, **p<0.01 (ANOVA with Tukey’s multiple comparison test comparing all groups). **(C)** Tumor size in area (mm^2^) of MCA205 stably expressing the globin-SL8-intron construct (left panel) and MCA205 WT (right panel) subcutaneously inoculated into the flank of immunocompetent C57BL/6 mice thereafter treated intraperitoneally with 30mg/kg of IP2 every three days once tumor size had reached 25-30mm^2^. Data are presented at the day when the ethical endpoints are reached and given as mean ± SEM. *P<0.05, **P<0.01, ***P<0.001 (unpaired student t test).

**Supplementary Fig. S8: The types of peptides displayed on H-2Kb and H-2Db molecules are altered by IP2.**

Length of the peptides found exclusively on treated or exclusively on control **(A)** MCA205 sarcoma cells and **(B)** B16F10 melanoma cells. Predicted MHC binding affinity of peptides found on **(C)** MCA205 sarcoma cells and **(D)** B16F10 melanoma cells computed on the NetMHC 4.0 Server. Peptides with an IC_50_<500nM are referred as strong binders. Data are given as mean ± SEM. *P<0.05, **P<0.01, ***P<0.001 (unpaired student t test).

