## Supplementary figures and images for "Splicing inhibition enhances the antitumor immune response through increased tumor antigen presentation and altered MHC-I immunopeptidome"

### Supplemental Figures

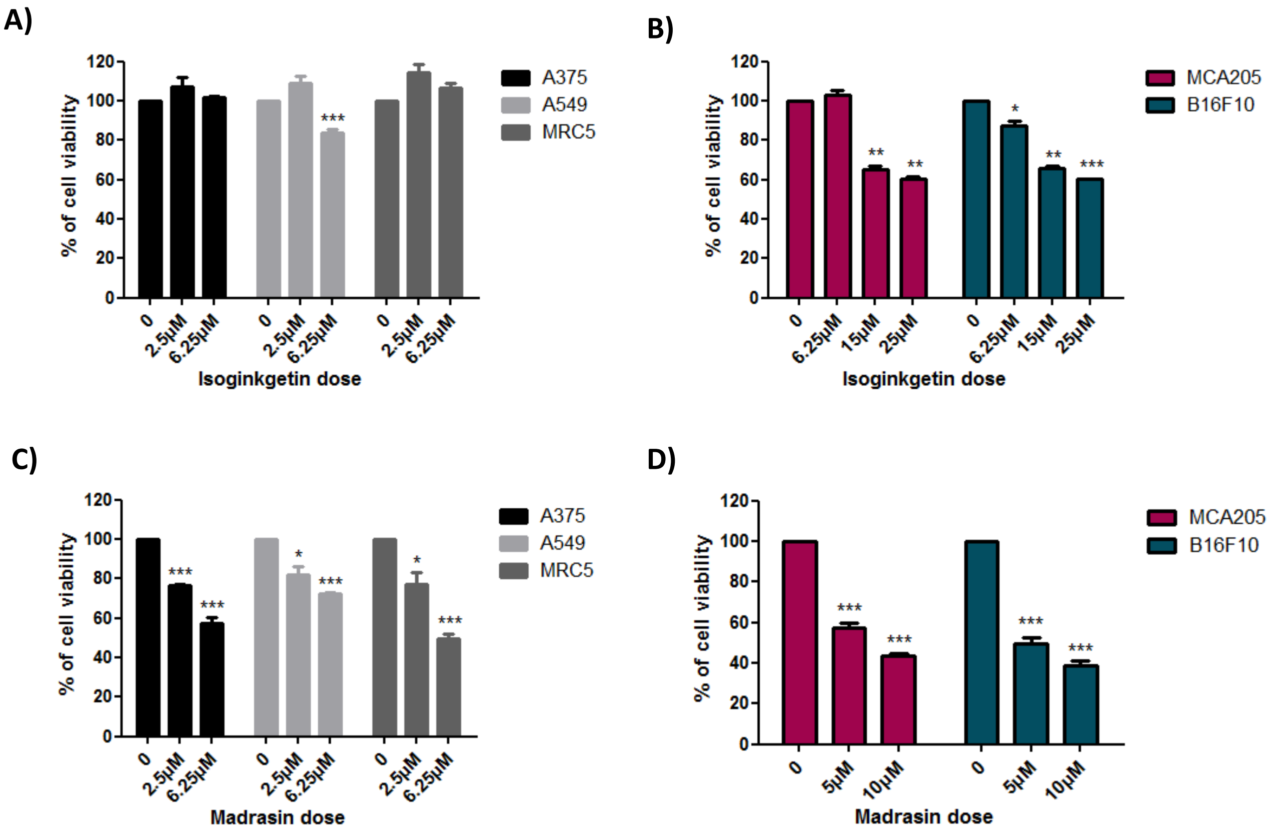

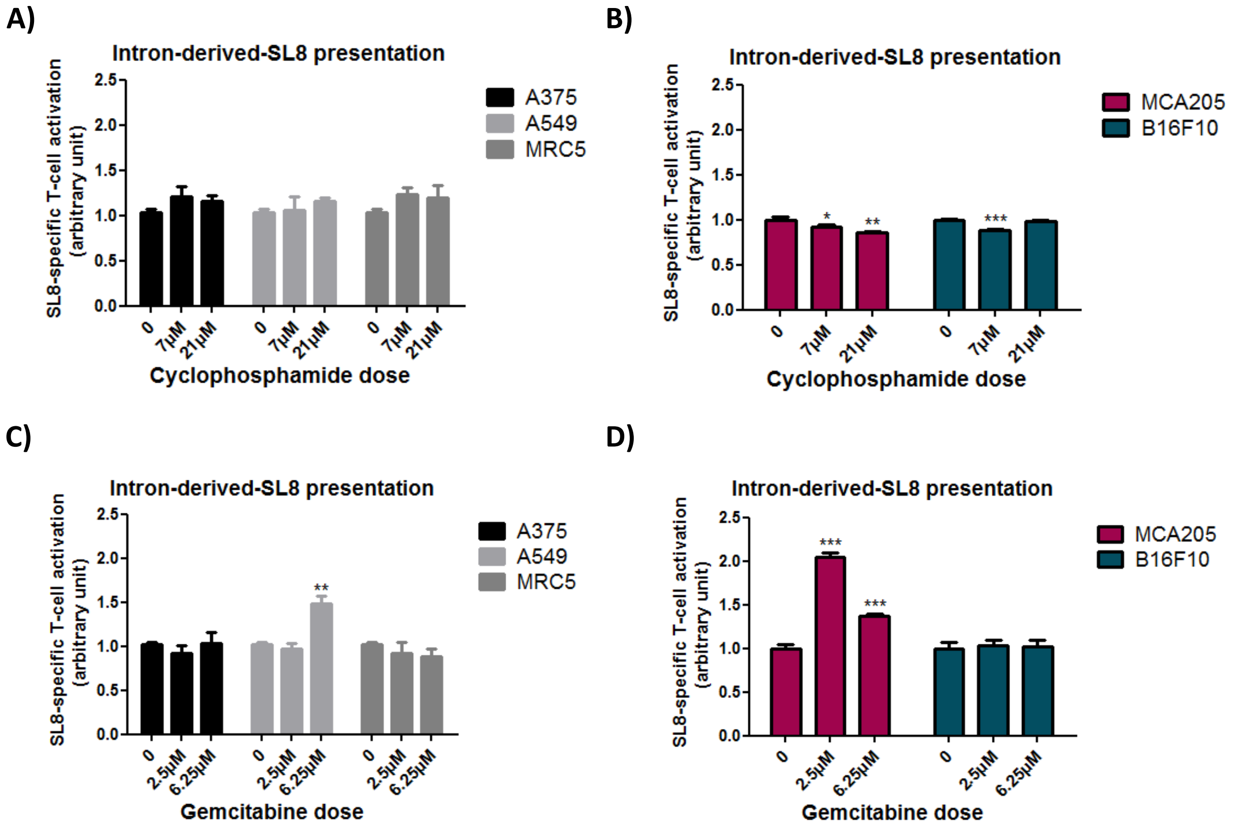

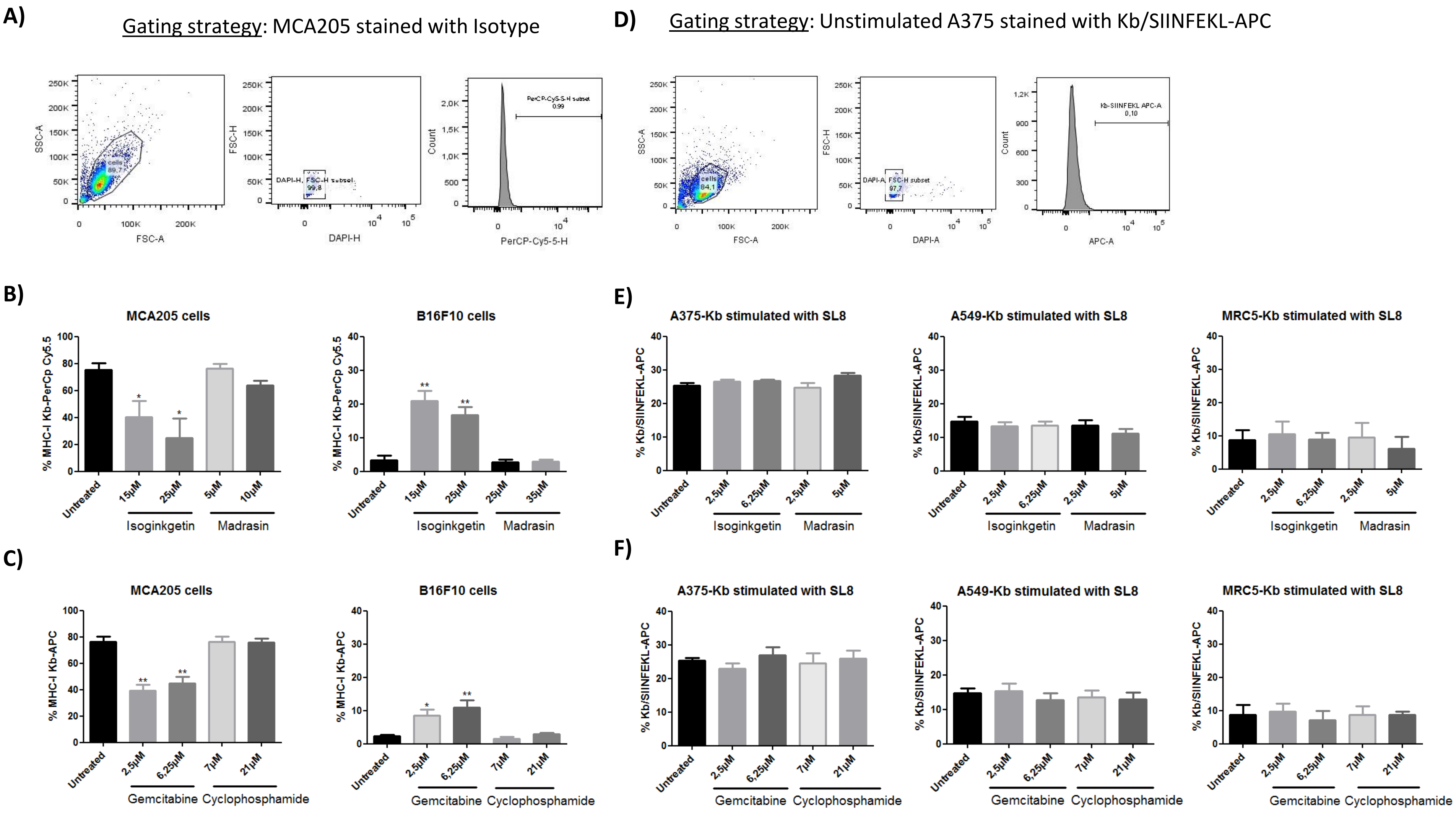

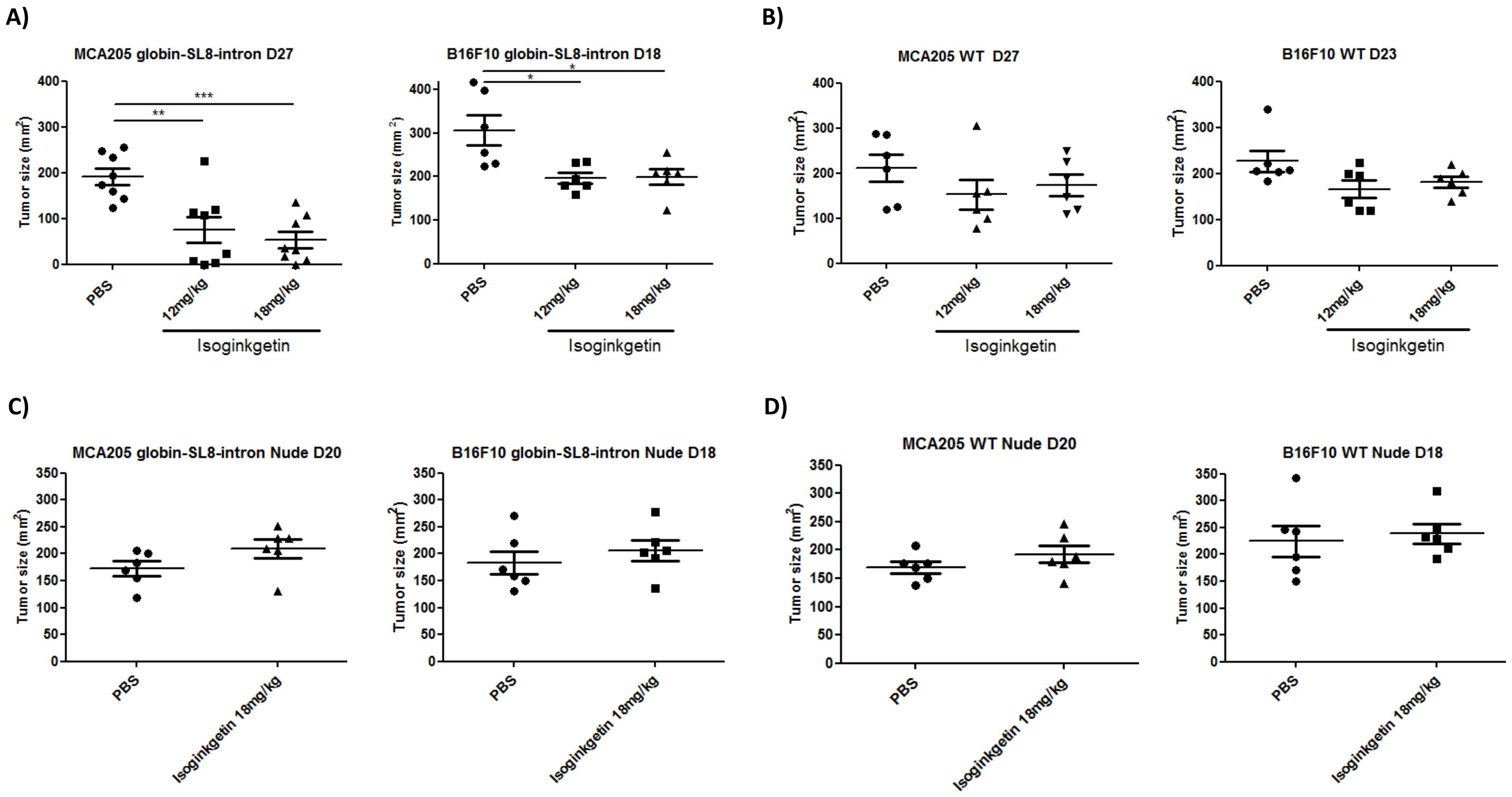

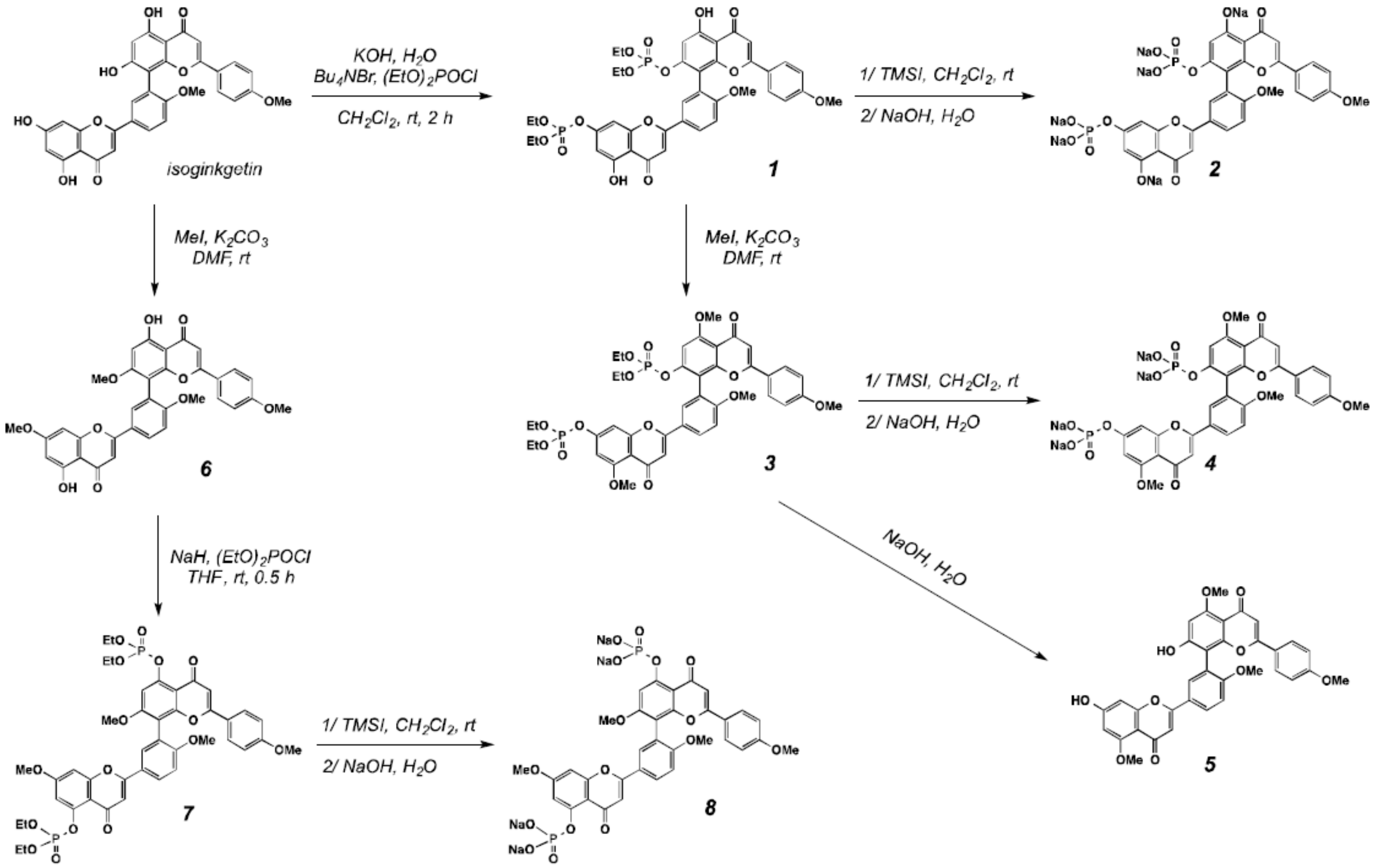

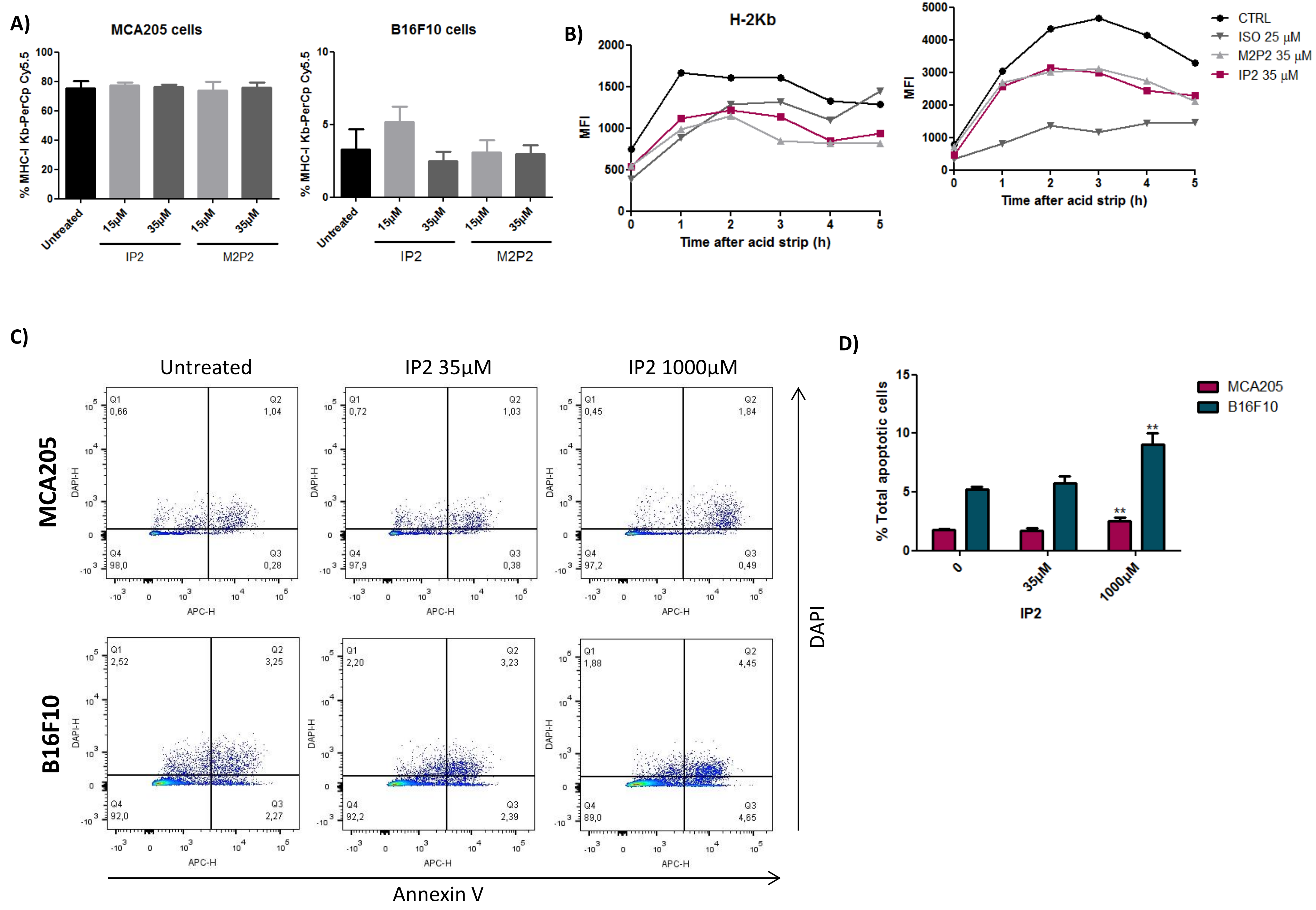

#### Figure S7

A)

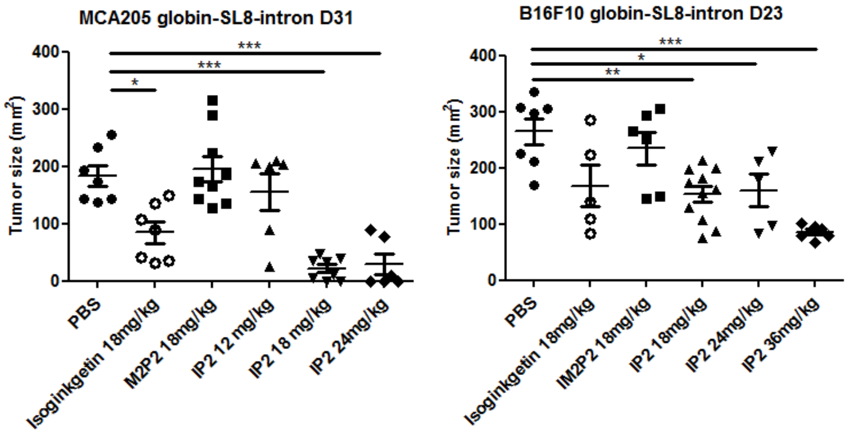


B)

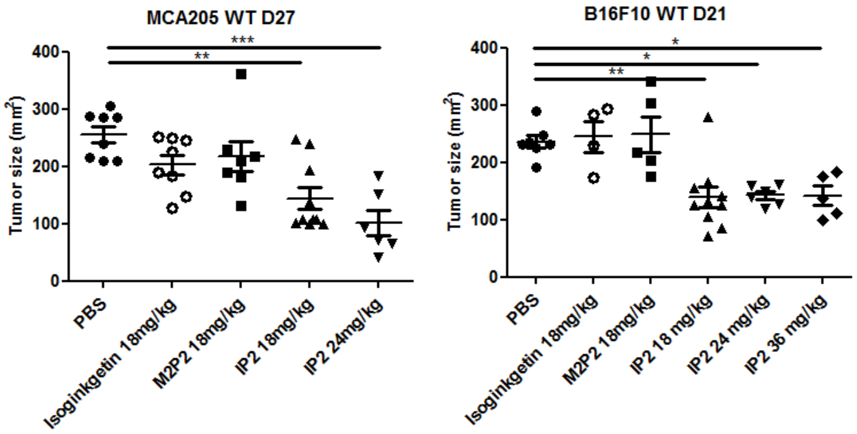


C)

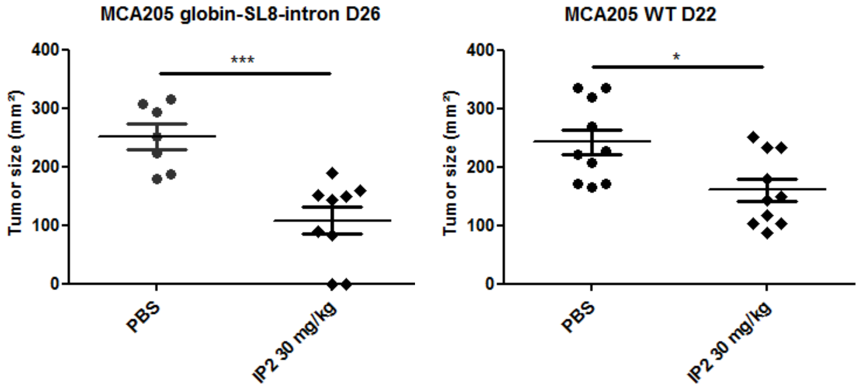

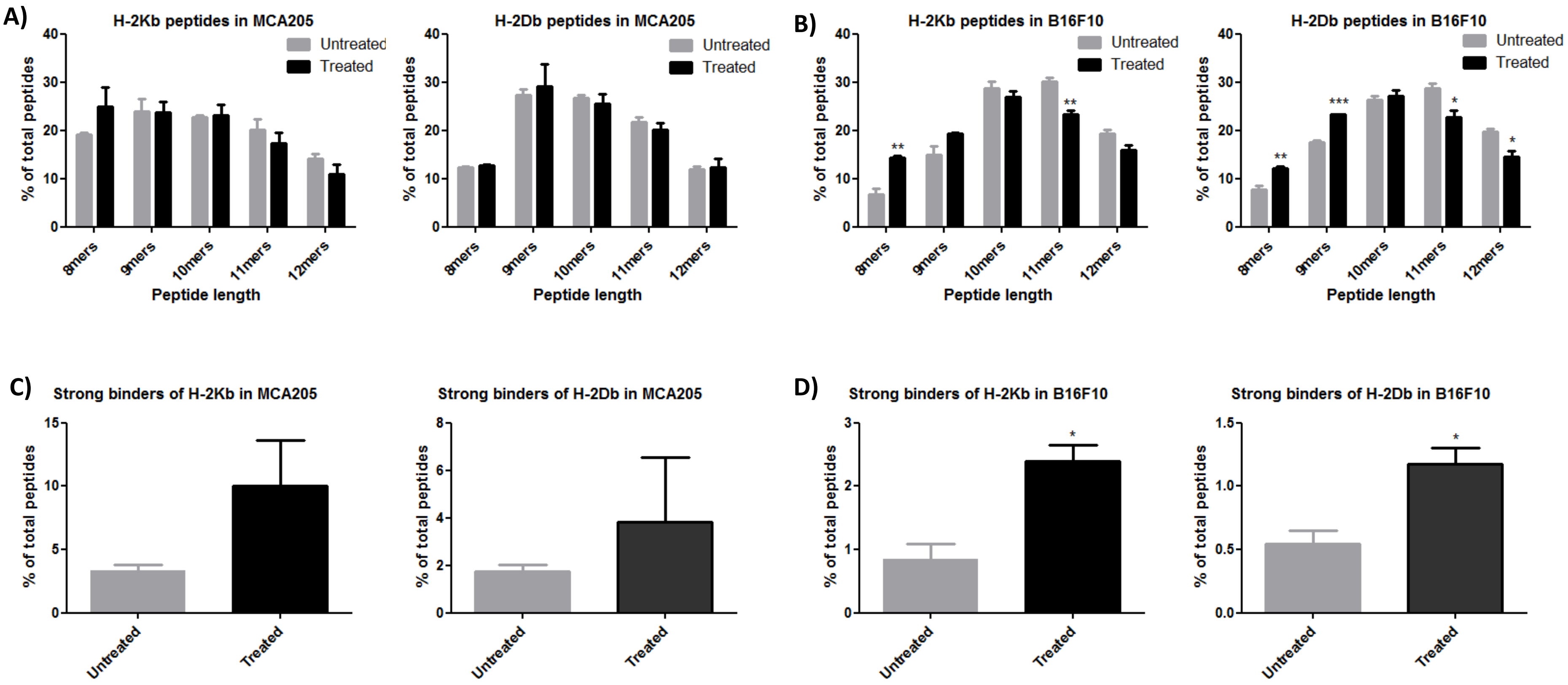
